# Attenuation of entorhinal cortex hyperactivity reduces Aβ and tau pathology

**DOI:** 10.1101/487405

**Authors:** Gustavo A Rodriguez, Geoffrey M Barrett, Karen E Duff, S. Abid Hussaini

## Abstract

High levels of the amyloid-beta (Aβ) peptide have been shown to disrupt neuronal function and induce hyperexcitability, but it is unclear what effects Aβ-associated hyperexcitability may have on tauopathy pathogenesis or propagation *in vivo*. Using a novel transgenic mouse line to model the impact of hAPP/Aβ accumulation on tauopathy in the entorhinal cortex-hippocampal (EC-HIPP) network, we demonstrate that hAPP aggravates EC tau aggregation and accelerates pathological tau spread into the hippocampus. *In vivo* recordings revealed a strong role for hAPP/Aβ, but not tau, in the emergence of EC neuronal hyperactivity and impaired theta rhythmicity. Chronic chemogenetic attenuation of Aβ-associated hyperactivity led to reduced hAPP/Aβ accumulation and reduced pathological tau spread into downstream hippocampus. These data strongly support the hypothesis that in Alzheimer’s disease (AD), Aβ-associated hyperactivity accelerates the progression of pathological tau along vulnerable neuronal circuits, and demonstrates the utility of chronic, neuromodulatory approaches in ameliorating AD pathology *in vivo*.

## Introduction

The accumulation of hyperphosphorylated, misfolded tau proteins into neurofibrillary tangles (NFT), coupled with deposition of amyloid beta (Aβ) into extracellular plaques, are two hallmark pathological features of Alzheimer’s disease (AD) in the brain. The severity of cortical NFT accumulation is strongly correlated with Aβ plaque load (1, 2) and is the principal neuropathological variable associated with cognitive impairment in AD (1-4). The entorhinal cortex (EC; Brodmann Areas 28 and 34) is a structure in the parahippocampal gyrus that plays a critical role in spatial representation and navigation (5-7), and it is one of the first structures to exhibit AD-related tauopathy and subsequent neuronal loss (8, 9). As AD progresses, considerable accumulation of pathological tau continues downstream into the hippocampus (HIPP), which is extensively connected to the EC. Preclinical investigation into the stereotypical spread of pathological tau along neuronal circuits in AD is an active area of research interest (10-13). However, the biological mechanisms underlying the propagation of tau pathology in the brain are currently unresolved.

*In vitro* studies utilizing rodent primary neurons have provided several mechanistic insights into the pathophysiological relationship between cleaved amyloid precursor protein (APP) fragments and tau. At the cellular level, accumulating evidence implicates Aβ oligomers as a causative agent in the increased phosphorylation of tau at AD relevant epitopes (14) and the missorting of tau and neurofilaments within the cell (15). In mouse models of tauopathy, stereotaxic injection of Aβ oligomers and fibrils into the brain results in significantly elevated phosphorylation of tau (16) and the increased induction of neurofibrillary tangles (NFT) (17). Thus, tauopathy in the brain may be aggravated by the increased production or accumulation of APP fragments *in vivo* through direct interaction. For reviews of experimental models that examine Aβ-induced tau alterations and pathology, see (18, 19).

Alternatively, human APP/Aβ accumulation in the brain may trigger the aggregation and acceleration of tau pathology via an intermediate, non-pathogenic mechanism. Indeed, several reports now describe an effect of Aβ accumulation on neuronal network hyperactivity in Aβ generating mouse models (20-22); for review, see (23, 24), as well as in humans with mild cognitive impairment (25, 26). Spontaneous, non-convulsive epileptiform-like activity has been described in cortical and hippocampal networks of relatively young transgenic mutant hAPP-expressing mice (21). In addition, increased proportions of neurons surrounding amyloid plaques exhibit aberrant hyperactivity (20) and are accompanied by the breakdown of slow-wave oscillations (27). Interestingly, neuronal hyperactivity has been shown to precede amyloid plaque formation in the hippocampus, suggesting that the abnormal accumulation of soluble Aβ drives aberrant neuronal network activity (28, 29). Thus, it is plausible that Aβ-associated hyperactivity can facilitate the progression of pathological tau in the brain, and does so indirectly without the need for direct Aβ-tau interaction. Interestingly, stimulating neuronal activity can facilitate both Aβ and tau release from neurons *in vivo* (30, 31), and exacerbates Aβ deposition and tauopathy in synaptically connected neurons (13, 32-34). Mature tau pathology may in turn aggravate Aβ-associated neuronal network dysfunction by further altering neuronal firing rates and network oscillations (35), recruiting neuronal populations into a harmful feedback loop involving protein aggregation and aberrant signaling.

In these studies, we utilize a newly developed AD mouse line to resolve the individual effects of Aβ and tau pathology on neuronal activity in the EC. Mice that generate Aβ and tau pathology were compared to littermates that generate either Aβ or tau alone, while non-transgenic littermates served as controls. We first demonstrate that hAPP/Aβ expression aggravates tau accumulation in the EC and accelerates pathological tau spread into the HIPP, supporting previous findings (36-38). *In vivo* electrophysiological recordings in our mice revealed distinct neuronal hyperactivity and network dysfunction associated with EC hAPP/Aβ expression, but not tau expression. We then employed a chemogenetic approach in the transgenic mice to combat EC neuronal hyperactivity, with the goal of reducing the accumulation of pathological Aβ and tau along the entorhinal cortex – hippocampal (EC-HIPP) network. Chronic attenuation of EC neuronal activity dramatically reduced hAPP/Aβ accumulation in downstream hippocampus and reduced abnormally conformed and hyper-phosphorylated tau aggregates along the EC-HIPP network. Our data support the emerging view that Aβ-associated hyperactivity plays a role in AD pathogenesis, specifically by acting as an accelerant of tau spread along synaptically connected neuronal circuits in the brain.

## Results

### Aβ-associated acceleration of tau pathology in EC-Tau/hAPP mice *in vivo*

Overexpression of mutant hAPP _(Swedish/Indiana)_ in the hAPP/J20 mouse line leads to progressive Aβ plaque deposition throughout the hippocampus and neocortex, contributing to aberrant network activity and cognitive deficits (21, 22). To test whether Aβ pathology alters the progression of tau pathology along the EC-HIPP circuit, we generated the hAPP/J20 x EC-Tau mouse line, hereafter referred to as EC-Tau/hAPP (described in Methods). At 16-months, EC-Tau/hAPP mice exhibit robust amyloid deposition throughout the EC and HIPP using 6E10 (Figure 1B), with no change in Aβ deposition compared to age-matched hAPP mice sampled. Diffuse amyloid accumulation made up the majority of the pathology, though small, compact Aβ plaques and large, dense-core Aβ plaques were also present (arrows, Figure 1B). Mice expressing hTau alone (EC-Tau) did not exhibit 6E10 immunoreactivity (Figure 1A). Immunostaining for MC1 revealed a dramatic increase in abnormal, misfolded tau within the somatodendritic compartments of EC and HIPP neurons of EC-Tau/hAPP mice compared to EC-Tau littermates (Figure 1C-D). This increase was over three-fold higher in EC cells (EC-Tau/hAPP: 131.00 ± 19.38 vs EC-Tau: 40.52 ± 7.24) and over ten-fold higher in DG granule cells (EC-Tau/hAPP: 359.20 ± 109.40 vs EC-Tau: 30.67 ± 14.88) (Figure 1E), suggesting that hAPP/Aβ expression in the EC-HIPP network accelerates tau pathology along the classical perforant pathway. MC1 immunostaining in 10-month EC-Tau/hAPP mice revealed mostly diffuse tau in neuropil throughout the EC and in axons terminating in the middle- and outer-molecular layers of the DG (Figure 1F). No somatodendritic MC1+ staining was detected in HIPP subregions at this age. By 16-months, it was clear that tau aggregation had not only increased in the EC, but had also spread into the HIPP. This accumulation of misfolded tau in the HIPP was notably heightened by 23-months. Thus, we chose to examine EC neuronal activity patterns in 16-month EC-Tau/hAPP mice to determine whether hAPP/Aβ was negatively impacting the local neuronal population and driving tau pathology.

**Figure 1.**
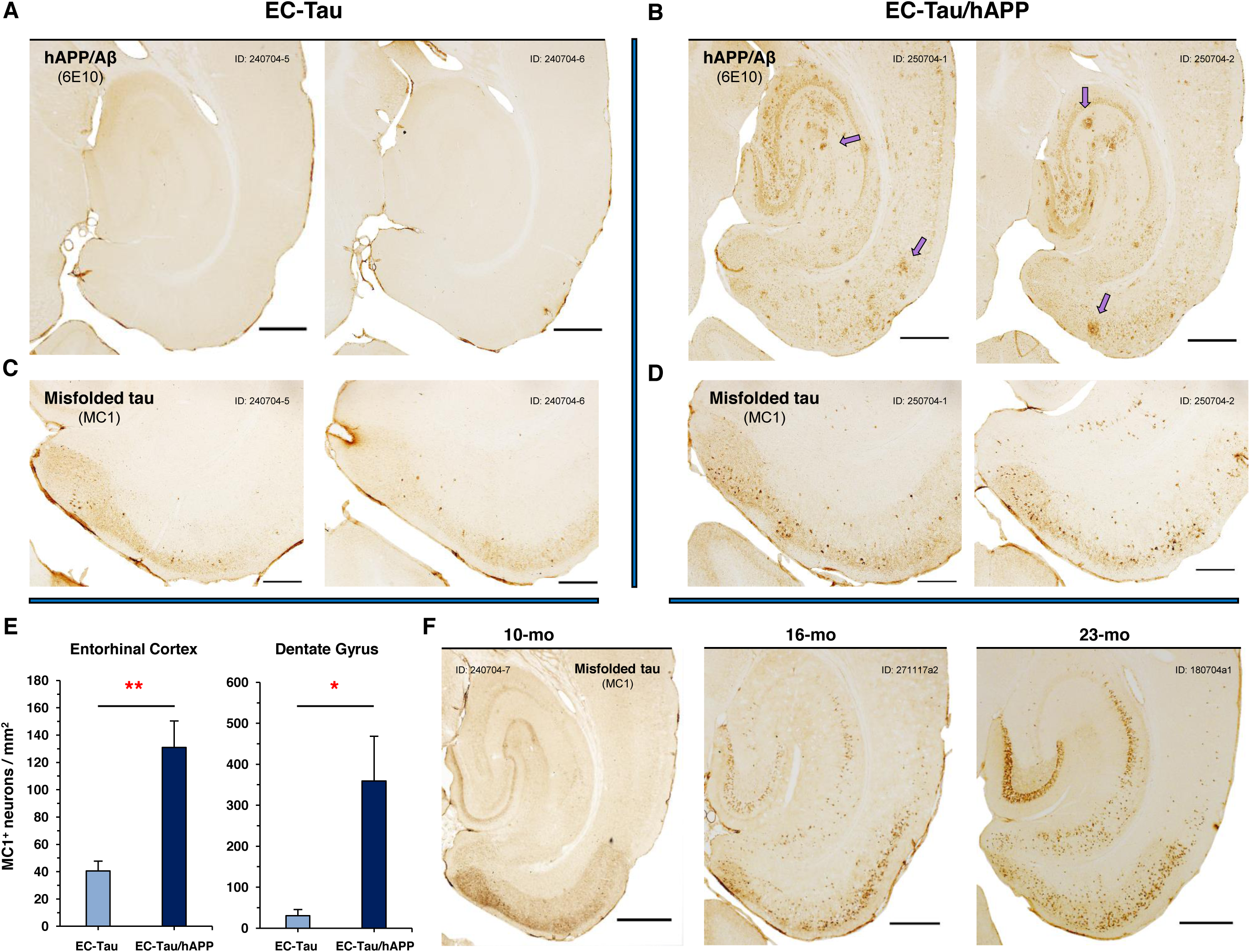
Aβ-associated acceleration of tau pathology along the EC-HIPP network. Horizontal brain sections from EC-Tau/hAPP mice and age-matched EC-Tau littermates were processed for immunohistochemical detection of hAPP/Aβ (6E10) and abnormal, misfolded tau (MC1). Representative, adjacent brain sections from two mice sampled are shown for both 6E10 and MC1. ***A-B***. 16-month EC-Tau/hAPP mice exhibited robust Aβ accumulation and plaque deposition throughout the EC and HIPP. Diffuse Aβ accumulation comprised the majority of the pathology in these regions, with occasional small, compact plaques and large, dense-core Aβ plaques present (*arrows*). EC-Tau mice did not exhibit 6E10+ immunoreactivity. Scale bars, 500µm. ***C-D***. MC1+ immunostaining revealed a clear acceleration of tau pathology in the EC of EC-Tau/hAPP mice, characterized by an increased number of cells with abnormally conformed tau localized to the somatodendritic compartment. Scale bars, 250µm. ***E***. A semiquantitative analysis of MC1+ cell counts was performed in the EC and DG. EC-Tau/hAPP brain sections exhibited over three-fold and over ten-fold greater MC1+ cell counts in EC and DG than in EC-Tau sections, respectively. Unpaired t-tests with Welch’s correction: EC, *p* < 0.01; DG, *p* < 0.05. ***F***. MC1+ immunostaining in EC-Tau/hAPP mice sampled at 10-, 16-, and 23-months of age confirmed the onset of pathological tau spread from the EC into the HIPP at 16-months. Graphs represent mean ± SEM for the averaged ROI values from three independently processed brain sections per mouse. * *p* < 0.05; ** *p* < 0.01.

### Aβ-associated EC hyperactivity and network dysfunction

*In vivo* multi-electrode recordings were performed in the EC of 16-month EC-Tau/hAPP mice and their age-matched, transgenic and non-transgenic littermates, as previously described (39). Briefly, tetrodes from custom-built microdrives (Axona, UK) were positioned to target cell layers II/III in the dorsal EC (see methods). Single-unit activity and local field potentials (LFPs) were collected while mice freely explored a large open field arena. Each experimental mouse underwent four to six recording sessions total over the course of several days, with only one recording session performed per day. Tetrodes for each mouse were moved down 0.1 mm from their previous position 24hr prior to the next recording session, allowing stable electrode positioning and a robust sampling of EC LII/III neuronal activity for each mouse. IHC was performed on post-mortem brain sections at the end of the study to confirm electrode placement (Figure 2A) and to examine the distribution of Aβ and tau pathology in the EC-HIPP network.

**Figure 2.**
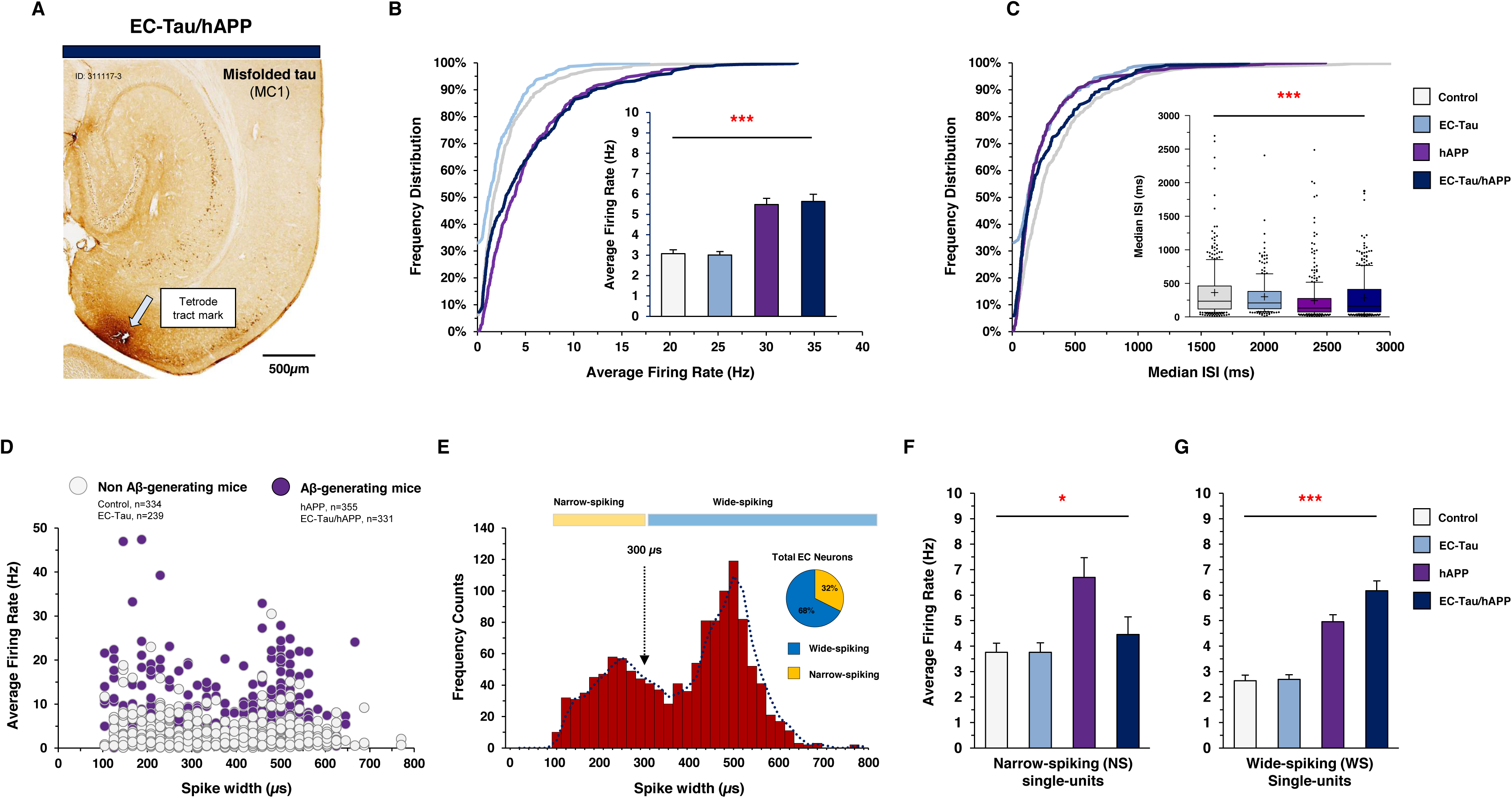
Aβ-associated hyperactivity in EC single-units *in vivo*. *In vivo* multi-electrode recordings were performed in the EC of 16-month EC-Tau/hAPP mice (n=4) and their age-matched littermates: EC-Tau (n=4), hAPP (n=5), and non-transgenic control mice (n=7). ***A.*** MC1 immunostained horizontal brain section from a 16-month EC-Tau/hAPP mouse. Representative image depicts tetrode tract mark terminating in EC layer II. Scale bar, 500µm. ***B.*** Cumulative frequency distributions of the spontaneous, average firing rates of all single-units collected in EC-Tau/hAPP mice and age-matched littermates. Pooled EC-Tau/hAPP neurons and hAPP neurons each exhibit shifts in their distributions towards higher firing rates versus Control. Two-sample Kolmogorov-Smirnov test: (EC-Tau/hAPP vs Control: *p* < 0.001; hAPP vs Control: *p* < 0.001). *Insert*, histograms of mean average firing rates are shown for each genotype. The single-unit average firing rates of EC-Tau/hAPP mice and hAPP mice were nearly two-fold higher than Control. Kruskal-Wallis test: *p* < 0.001. ***C***. Cumulative frequency distributions of the median inter-spike intervals (ISI) for pooled EC single-units. Two-sample Kolmogorov-Smirnov test: (EC-Tau/hAPP vs Control: *p* < 0.001; hAPP vs Control: *p* < 0.001). *Insert*, box-and-whisker plots depicting median and mean ISI values along with 10-90 percentile limits per genotype. EC single-unit ISI values were significantly decreased in EC-Tau/hAPP mice and hAPP mice compared to Control. Kruskal-Wallis test: *p* < 0.001. ***D***. EC single-unit average firing rates were plotted as a function of waveform spike-width and color-coded based on whether they belonged to Aβ-generating mice (purple) or Non Aβ-generating mice (gray). A distinct population of hyperactive EC neurons were present in Aβ-generating mice, and were distributed along the x-axis (shorter to longer spike-widths). ***E.*** Frequency histogram of neuronal spike widths for all single-units; note the bimodal distribution. A cutoff of 300µm spike-width was used to delineate putative interneurons (NS, narrow-spiking cells; yellow) from putative excitatory neurons (WS, wide-spiking cells; blue) (*broken arrow*). ***F.*** hAPP NS cells exhibit increased average firing rates compared to Controls. Kruskal-Wallis test: *p* < 0.05. ***G***. EC-Tau/hAPP WS cells and hAPP WS cells each exhibit increased average firing rates versus Control WS cells. Kruskal-Wallis test: *p* < 0.001. Bar graphs represent mean ± SEM. * *p* < 0.05; *** *p* < 0.001.

A total of 1,260 EC neurons were recorded and analyzed from 20 mice (Control, n=7; EC-Tau, n=4; hAPP, n=5; EC-Tau/hAPP, n=4) (Figure 2) (for detailed methodology, see Methods). Plotting the cumulative frequency distributions of the spontaneous, average firing rates of all recorded neurons showed a clear shift towards higher firing rates in both EC-Tau/hAPP mice and hAPP mice versus Control mice (Figure 2B). Individual two-sample Kolmogorov-Smirnov tests comparing the distributions of EC-Tau/hAPP and hAPP neuronal firing rates to Controls confirmed this shift (*p* < 0.001). The average firing rates for pooled EC neurons in each genotype were then compared (Figure 2B, insert). EC-Tau/hAPP and hAPP neurons each exhibited nearly a two-fold increase in their average firing rates versus Control neurons. Interestingly, the average firing rates of EC-Tau neurons was not significantly different from Controls, suggesting that the increased hyperactivity in EC-Tau/hAPP neurons was driven by hAPP expression and/or Aβ deposition, but not tau. To examine task-relevant neuronal firing rates during active exploration, we applied a minimum speed filter (<5cm/sec) to the data to remove EC spiking activity occurring during bouts of immobility (Supplemental Figure 1A). hAPP (2.291 ± 0.141 Hz) and EC-Tau/hAPP (2.658 ± 0.170 Hz) neurons once again exhibited significantly increased average firing rates compared to Control (1.299 ± 0.092 Hz) in speed filtered datasets. We then examined the inter-spike interval (ISI) values for EC single-units as an additional measure of neuronal hyperactivity. Cumulative frequency distributions of the median ISIs showed that EC-Tau/hAPP and hAPP single-units were skewed towards shorter ISIs than Control cells for ∼65% of the data (Figure 2C). The distribution of median ISIs then begin to shift towards higher values in EC-Tau/hAPP mice (>70%) and group well with the remaining Control data. Interestingly, the median ISI values for EC-Tau cells were skewed towards shorter median ISIs. However, EC single-unit median ISI values were only significantly shorter in EC-Tau/hAPP mice (286.80 ± 16.86 ISI) and hAPP mice (242.40 ± 16.92 ISI) compared to Controls (363.10 ± 21.06 ISI). EC-Tau single-unit median ISI values (300.10 ± 18.08 ISI) were not significantly different from Control.

Interneuron dysfunction has previously been described in hAPP expressing mice, suggesting that shifts in the excitation-inhibition ratio within cortical and hippocampal networks may be responsible for emergent epileptiform activity (21, 40). We first plotted the average firing rates of all EC single-units as a function of the waveform’s averaged spike-width (µs). This revealed a distinct population of hyperactive neurons in Aβ-generating mice that were distributed along narrow and wide spike widths (Figure 2D, *purple*). We also noticed what appeared to be a bimodal distribution in the scatterplot. To examine cell-type specific firing patterns in our single-unit recordings, we then plotted cell frequency as a function of average spike width (Figure 2E) (39, 41). This resulted in a clear bimodal distribution of the cell population, wherein narrow-spiking (NS) cells could be separated from wide-spiking (WS) cells (Figure 2D-G). Previous studies indicate that NS cells and WS cells likely correspond to putative interneurons and putative excitatory cells, respectively (41). A total of 408 NS cells and 852 WS cells were identified from our EC recordings across 20 mice (NS, 32.38% vs WS, 67.62% of total neurons) (for detailed per animal cell-type information, see Table 1 and Methods). hAPP (6.694 ± 0.778 Hz) NS cells exhibited increased firing rates compared to Control (3.757 ± 0.358 Hz) NS cells, while EC-Tau/hAPP (4.456 ± 0.688 Hz) NS cells and EC-Tau (3.760 ± 0.367 Hz) NS cells did not (Figure 2F). Interestingly, hAPP (4.954 ± 0.275 Hz) WS cells and EC-Tau/hAPP (6.174 ± 0.388 Hz) WS cells exhibited significantly increased average firing rates compared to Control (2.693 ± 0.218 Hz) WS cells (Figure 2G). The average firing rate of EC-Tau (2.693 ± 0.180 Hz) WS cells did not differ from Controls. Applying a speed filter to the datasets revealed similar effects on EC firing rates (Supplemental Figure 1B-C). These data agree with previous reports describing Aβ-associated dysfunction in hippocampal interneurons, and expand this hyperactivity phenotype to both putative interneurons and putative excitatory neurons in EC.

**Table 1.** Single-unit totals and average firing rates per mouse. *In vivo* electrophysiological recording parameters are shown for individual mice included in the study. The total number of cells and their average firing rates are listed, as well the average firing rates and total numbers of narrow-spiking (NS) and wide-spiking (WS) cells per mouse. Finally, the averaged bursting activity % per mouse is listed.

|  | Control |  |  |  |  |  |  | EC-Tau |  |  |  | hAPP |  |  |  |  | EC-Tau/hAPP |  |  |  |
| --- | --- | --- | --- | --- | --- | --- | --- | --- | --- | --- | --- | --- | --- | --- | --- | --- | --- | --- | --- | --- |
|  | 270704a3 | 270704-3 | 290704-1 | 430704-1 | 430704-6 | 430704-7 | 450704-4 | 27805a4 | 306805a3 | 190805-5 | 195805a4 | 261031a3 | 301031a2 | 421031-1 | 421031-2 | 421031-5 | 271117a2 | 291117a2 | 311117-3 | 311117-4 |
| Total Cell # | 45 | 37 | 77 | 47 | 34 | 36 | 58 | 60 | 31 | 70 | 78 | 80 | 60 | 62 | 82 | 71 | 55 | 122 | 79 | 75 |
| Total Cell, AVG FR (HZ) | 3.177 | 1.924 | 2.835 | 3.117 | 2.573 | 3.655 | 4.084 | 3.468 | 2.84 | 2.614 | 3.066 | 6.161 | 5.429 | 4.217 | 5.75 | 5.54 | 4.617 | 7.452 | 3.72 | 5.462 |
| NS Cell, Total # | 12 | 7 | 40 | 28 | 5 | 15 | 19 | 21 | 1 | 24 | 24 | 11 | 10 | 14 | 49 | 25 | 44 | 21 | 31 | 7 |
| NS Cell, AVG FR (Hz) | 2.634 | 2.980 | 3.342 | 3.851 | 2.551 | 5.514 | 4.420 | 4.406 | 2.966 | 3.455 | 3.533 | 6.621 | 3.084 | 3.626 | 7.371 | 8.560 | 5.182 | 1.382 | 5.981 | 2.359 |
| WS Cell, Total # | 33 | 30 | 37 | 19 | 29 | 21 | 39 | 39 | 30 | 46 | 54 | 69 | 50 | 48 | 33 | 46 | 11 | 101 | 48 | 68 |
| WS Cell, AVG FR (Hz) | 1.665 | 3.223 | 2.287 | 2.036 | 2.576 | 2.328 | 3.920 | 2.963 | 2.836 | 2.175 | 2.859 | 6.088 | 5.898 | 4.390 | 3.449 | 3.895 | 2.355 | 8.715 | 2.259 | 5.781 |
| Bursting % | 3.717 | 0.914 | 3.158 | 2.754 | 1.604 | 3.338 | 2.503 | 3.233 | 2.800 | 1.533 | 1.120 | 3.931 | 3.264 | 2.514 | 4.144 | 4.755 | 3.878 | 4.810 | 2.312 | 2.633 |

Impaired neuronal network activity has been described in human AD (42) and in mouse models of AD pathology (43-45). To investigate the effects of hAPP/Aβ and tau pathology on EC network activity *in vivo*, we examined the LFPs and compared oscillatory activity across genotypes (Figure 3). Initial, visual inspection of the filtered LFP signal in theta (4-12 Hz), low gamma (35-55 Hz), and high gamma (65-120 Hz) frequency ranges, as well as the filtered LFP spectrograms, demonstrate impaired theta rhythmicity in 16-month EC-Tau/hAPP mice compared to non-transgenic Control mice (Figure 3A-B). The average % theta power values for EC-Tau/hAPP (24.85 ± 3.55 %) mice and hAPP (27.88 ± 2.51 %) mice were nearly one-half that of Control (43.50 ± 6.86 %) mice and EC-Tau (44.71 ± 4.29 %) mice. When quantified, EC-Tau/hAPP (n=8) mice and hAPP (n=9) mice exhibited a marked decrease in % power distribution within the theta frequency range compared to Control (n=7) mice. Theta power was not significantly affected in EC-Tau (n=5) mice (*p* > 0.05). No genotype differences were detected in the low gamma or high gamma frequency ranges. As running speed can impact theta power in the hippocampal formation (46, 47), we speed filtered our LFP recordings and re-analyzed the data to remove activity during bouts of immobility (Supplemental Figure 1D-F). Similarly, we found an Aβ-associated decrease in % theta power, though this was accompanied by an Aβ-associated increase in % low gamma power. Averaged, speed filtered % high gamma power values were not significantly different across genotypes.

**Figure 3.**
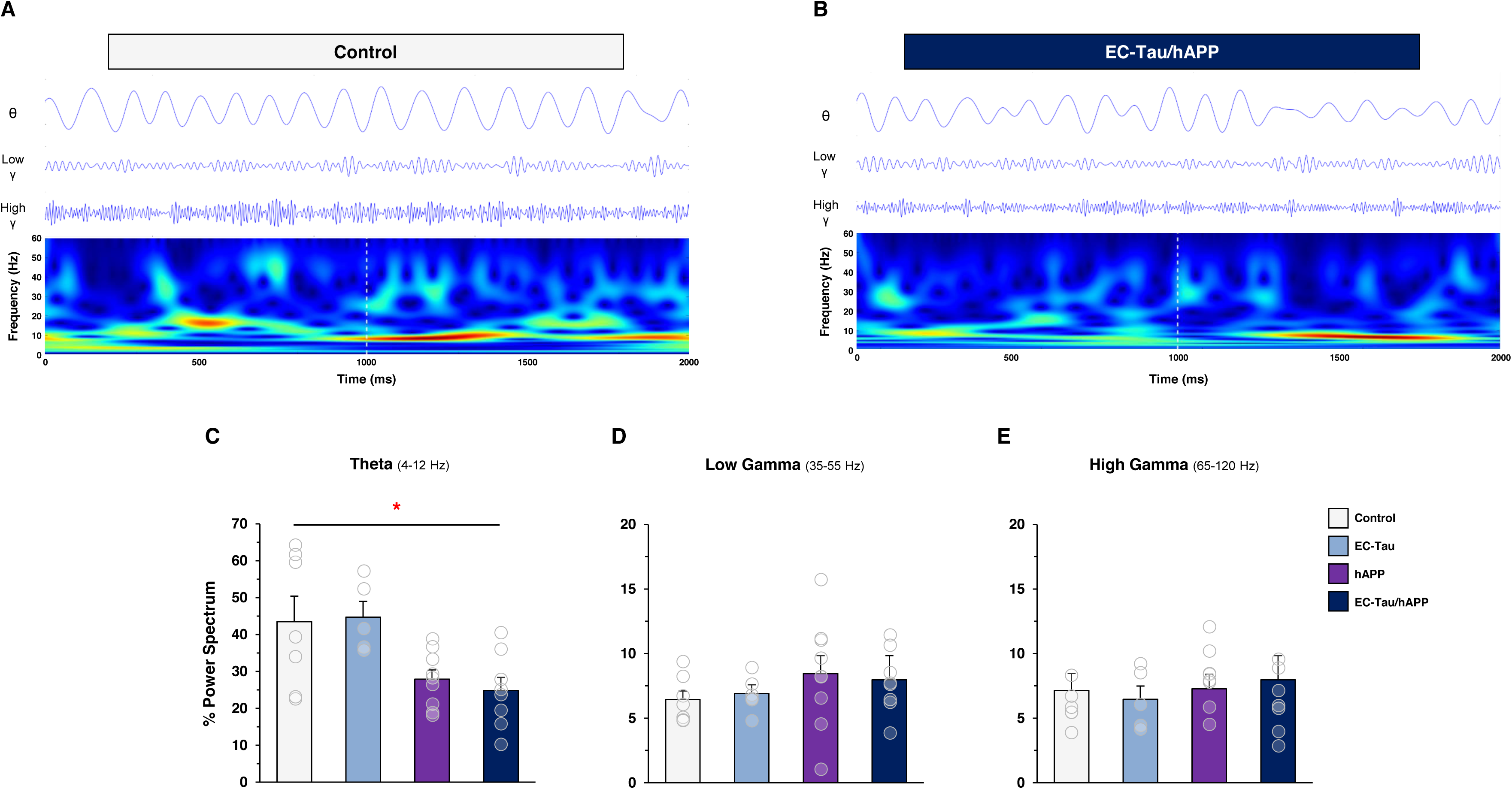
Aβ-associated impairment of EC network activity *in vivo*. Local field potentials (LFPs) were collected within the EC of 16-month EC-Tau/hAPP (n=8) mice and age-matched littermates (EC-Tau, n=5; hAPP, n=9; non-transgenic control, n=7). The percentage power values for oscillatory frequency bands were then calculated and compared across genotype. ***A-B.*** Representative, filtered LFP waveforms in the theta (4-12 Hz), low gamma (35-55 Hz), and high gamma (65-120 Hz) frequency ranges are shown for one Control mouse and one EC-Tau/hAPP mouse recording session, along with the LFP spectrograms. LFP signals show a clear disturbance in the theta rhythm of 16-month EC-Tau/hAPP mice. Time-scale, 2000 ms. ***C.*** Quantification of % power values revealed a significant decrease in averaged % theta power in EC-Tau/hAPP (24.85 ± 3.55 %) mice and hAPP (27.88 ± 2.51 %) mice versus Control (43.50 ± 6.86 %) mice. One-way ANOVA test: *p* < 0.01; Dunn’s multiple comparison test, *p* < 0.05. ***D-E.*** No differences in averaged % power spectrum values were detected across genotypes in the low gamma (*p* > 0.05) or high gamma (*p* > 0.05) frequency ranges. Bar graphs represent mean ± SEM. Transparent overlays represent individual % power spectrum values per mouse in each genotype. ** *p* < 0.01.

Overexpression of hAPP_Swedish/Indiana_ has previously been associated with increased locomotor activity in the hAPP/J20 mouse line (22, 48, 49). To investigate locomotor activity in 16-month EC-Tau/hAPP mice, we analyzed the position data from our recorded mice as they performed a foraging task in the open field arena (Figure 4). We did not detect significant differences between groups in the total distance traveled in the arena (m), % of arena coverage or average speed (cm/sec) during exploration (Figure 4A-D). To examine whether group differences in locomotor activity exist within the initial phases of recording, we split the sessions and examined performance measures in the first 5, 10 and 15 min (Figure 4E-G). No significant differences were detected between groups in any measure and in any time bin examined (*p* > 0.05), suggesting that Aβ and tau pathology do not affect motivated foraging behavior in an open field arena.

**Figure 4.**
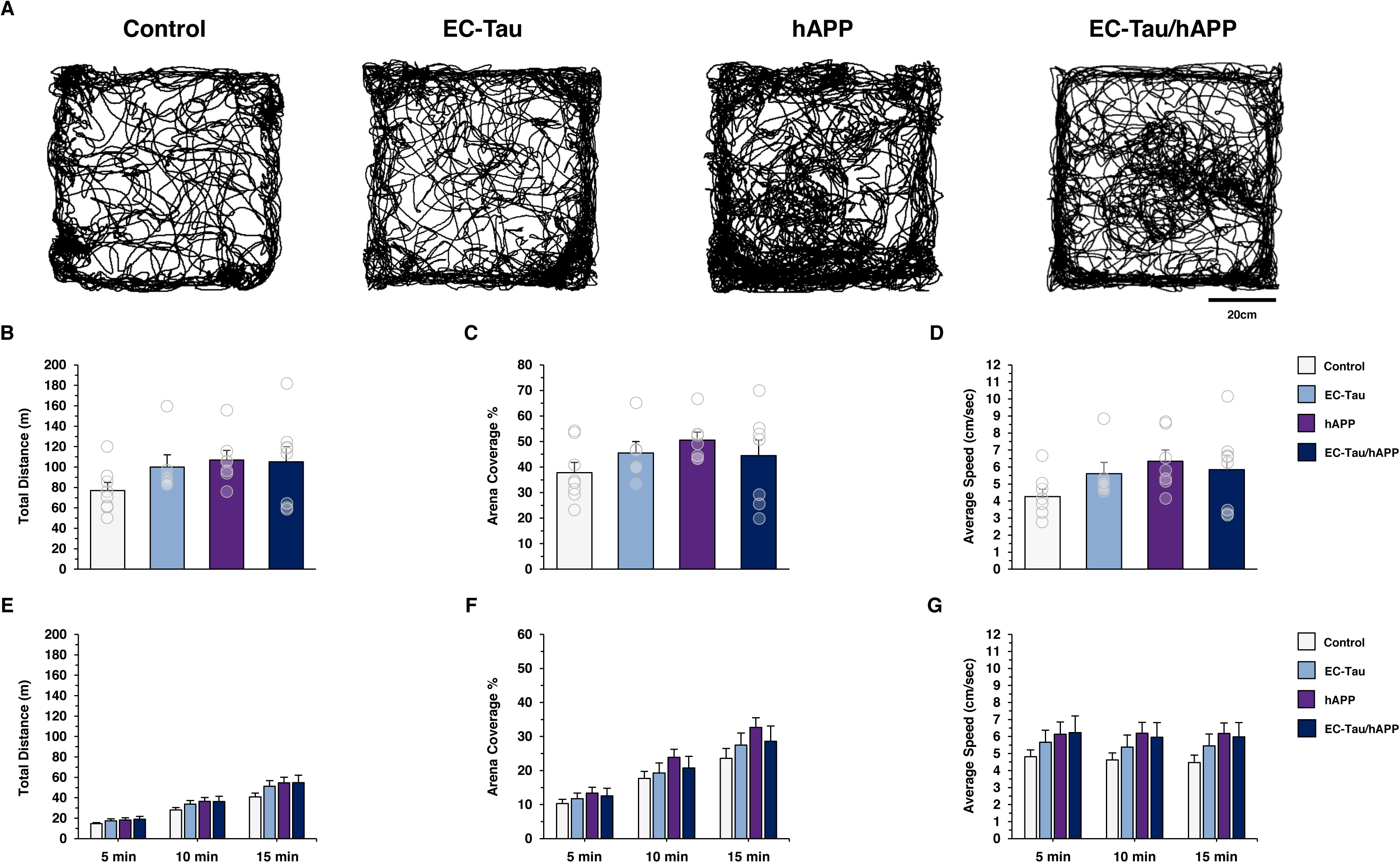
Aβ and tau pathology does not affect locomotor activity in an open field. Locomotor activity was assessed by analyzing the position data of each mouse during active exploration in an open field (recording session, 30 min). ***A***. Representative trajectories are shown for one recording session per genotype. Scale bar, 20cm. ***B-D***. The following parameters in the open field were analyzed and compared across genotype: the total distance traveled (m), the % arena coverage and average speed (cm/sec). No significant group differences were detected on any measure. One-way ANOVA tests: Total distance (m), F_(3,25)_ = 1.540, *p* > 0.05; % of arena coverage, F_(3,25)_ = 1.281, *p* > 0.05; average speed (cm/sec), F_(3,25)_ = 1.632, *p* > 0.05. ***E-G***. No significant group differences were detected on any dependent measure when analyzing position data in the first 5, 10 or 15 min of the recording sessions (One-way ANOVA tests; *p* > 0.05). Bar graphs represent mean ± SEM. Individual values (transparent overlays) are representative means from three averaged recording sessions per mouse.

Taken together, our *in vivo* recording data in the EC-Tau/hAPP mouse line suggests that at 16-months, hAPP/Aβ expression is associated with EC neuronal hyperactivity in both putative interneurons and excitatory neurons, and that tau expression may selectively blunt putative interneuron hyperactivity in EC-Tau/hAPP mice. hAPP/Aβ expression is also associated with a distinct decrease in % theta power in both speed filtered and unfiltered EC LFPs. Finally, tau pathology does not appear to overtly impact EC neuronal activity at 16-months.

### Chemogenetic attenuation of EC hyperactivity reduces Aβ accumulation in downstream hippocampus

Previous reports have shown that elevating neuronal activity can increase Aβ deposition and tau pathology *in vivo* (32-34). However, these effects were restricted to mouse lines that individually overexpress hAPP and deposit Aβ/amyloid, or overexpress mutant hTau and accumulate pathological tau in the brain. To date, no studies have examined the effects of Aβ-associated neuronal hyperactivity on tau pathology *in vivo*, or whether direct neuromodulatory intervention at the site of hyperactivity can impact the acceleration of tau pathology along neuronal networks *in vivo*. Therefore, we employed a chemogenetic approach in a series of experiments wherein a modified form of the human muscarinic M4 receptor was virally delivered and expressed in the EC of 16-month EC-Tau/hAPP mice and activated for 6-weeks. Activation of hM4D_i_ DREADD receptors (designer receptor exclusively activated by a designer drug) via clozapine-n-oxide (CNO) engages the G_i_ signaling pathway and effectively reduces neuronal firing rates in transduced cell populations (50, 51).

Single, high dose CNO (5 and 10mg/kg, i.p.) injections reliably induced DREADD expression and altered EC neuronal activity and theta power in EC-Tau mice and hAPP mice (Supplemental Figure 2), providing valuable metrics for detecting chronic DREADDs activation *in vivo*. Following EC recording experiments, 16-month EC-Tau/hAPP mice (n=4) and age-matched, transgenic littermates (EC-Tau, n=5; hAPP, n=3) expressing hM4D_i_ DREADDs targeted to EC excitatory neurons were implanted with osmotic minipumps loaded with CNO. hM4D_i_ EC DREADDS receptors were activated for 6-weeks by continuous delivery of CNO (1mg^-1^/kg^-1^/day^-1^) into the peritoneum. EC spiking activity was noticeably decreased during the last three weeks of CNO treatment (n=9 mice, EC-Tau/hAPP and EC-Tau mice) (Figure 5A), and theta power was significantly reduced at the fifth (T5) and sixth week (T6) of CNO treatment compared to baseline (Figure 5B), confirming chronic hM4D_i_ DREADDs activation *in vivo*. After 6-weeks of hM4D_i_ EC DREADDs activation, all experimental mice were sacrificed and immunostaining was performed on horizontal brain sections to confirm EC DREADDs expression and electrode placement, and to identify pathological Aβ and/or tau deposition along the EC-HIPP network. An overlay of mCherry signal (hM4D_i_ DREADDs) and eGFP (control virus) expression patterns for EC-Tau/hAPP mice (n=4) and transgenic littermates (hAPP, n=3; EC-Tau, n=5) is shown (Figure 5C & Supplemental Figure 4A-G). hM4D_i_ DREADDs expression in the right hemisphere was primarily localized to cell bodies and neuropil throughout the EC and pre- and para-subiculum, with occasional mCherry signal present in the subiculum, and at terminal ends of axons in the middle- and outer-molecular layers of the DG (Figure 5C-D). Importantly, we did not detect DREADDs mCherry crossover into the contralateral left hemisphere.

**Figure 5.**
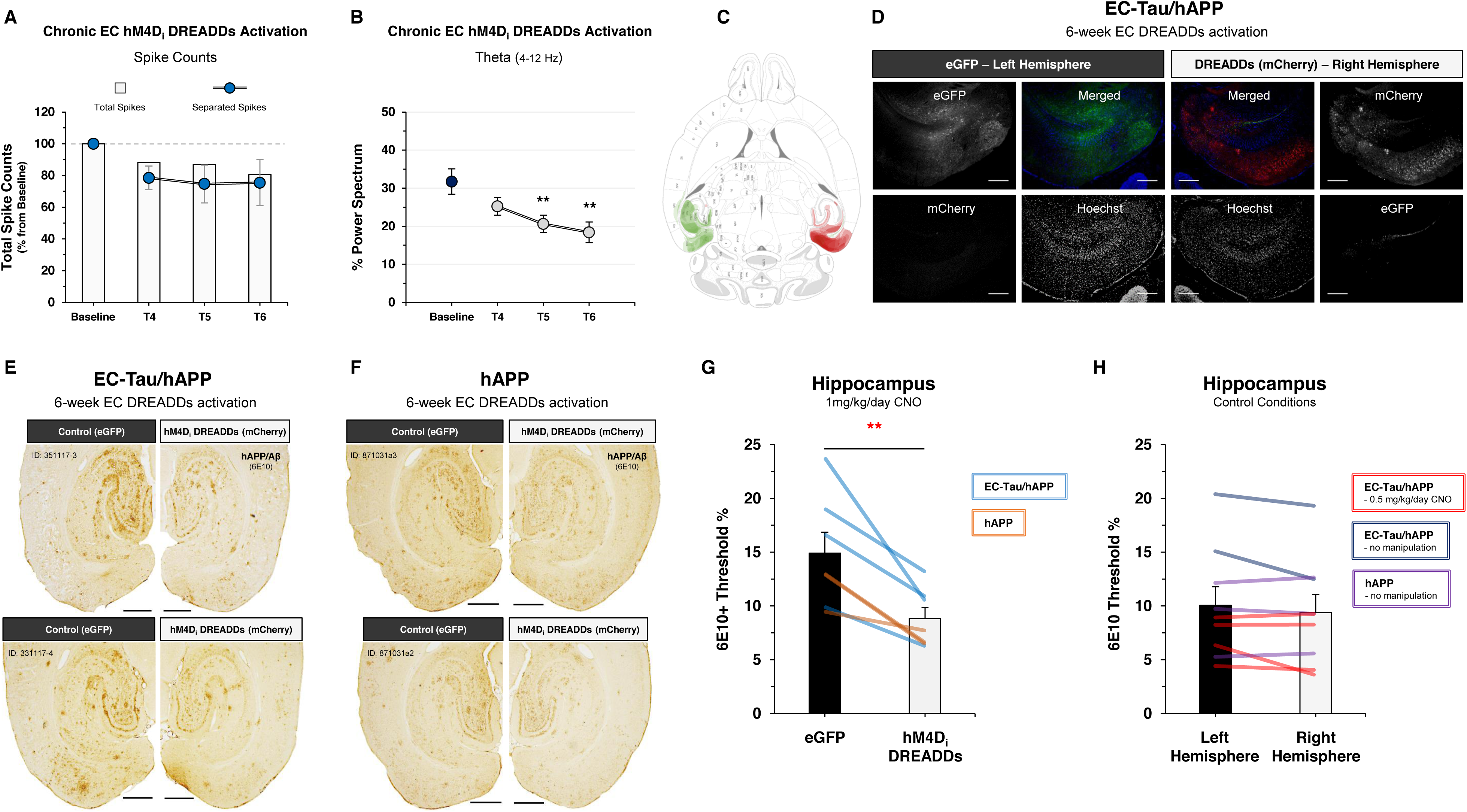
Chronic hM4D_i_ EC DREADDs activation reduces Aβ pathology in downstream hippocampus. 16-month EC-Tau/hAPP (n=4) mice and age-matched littermates (EC-Tau, n=5; hAPP, n=3) expressing hM4D_i_ EC DREADDS were subjected to 6-weeks of DREADDS activation via osmotic minipumps (CNO, 1mg^-1^/kg^-1^/day^-1^). ***A-B.*** Neuronal activity was examined *in vivo* for EC-Tau/hAPP and EC-Tau mice (n=9) and analyzed at time-points corresponding to the last three weeks of drug treatment (T4, T5 and T6). Chronic hM4D_i_ EC DREADDs activation reduced total spike counts from separated cells (∼25% reduction), and reduced % theta power versus baseline recordings. Repeated measures ANOVA: *F*_(3,7)_ = 5.157, *p* < 0.01. Dunnett’s multiple comparison test. ***C-D.*** An overlay of individual mCherry (hM4D_i_ DREADDs) and eGFP (control virus) expression patterns for EC-Tau/hAPP mice and transgenic littermates is shown (n = 12 mice). hM4D_i_ DREADDs expression in the right hemisphere was localized to cell bodies and neuropil throughout the EC, pre- and para-subiculum, and subiculum, with occasional mCherry signal present at terminal ends of axons in the outer 2/3 molecular layer of DG. Representative IF images from a 16-month EC-Tau/hAPP mouse are shown. Scale bars, 250 µm. ***E-F.*** 6E10+-immunostained brain sections are shown for two EC-Tau/hAPP and hAPP mice. hAPP/Aβ accumulation is reduced in ipsilateral HIPP compared to contralateral HIPP. Scale bars, 500µm. ***G.*** Semiquantitative analysis of HIPP 6E10+ immunoreactivity was performed and compared across hemispheres (EC-Tau/hAPP, n=4; hAPP, n=3). Image thresholding analysis revealed a significant reduction in 6E10+ immunoreactivity in the ipsilateral HIPP, expressed as a % of the HIPP region (ipsi HIPP: 8.85 ± 1.03 % vs contra HIPP: 14.92 ± 1.95 %). Paired *t*-test: *t* (6) = 4.552, *p* < 0.01. ***H.*** HIPP 6E10+ immunoreactivity did not differ between right and left hemispheres in 16-18 month EC-Tau/hAPP (n=2) mice and hAPP (n=4) mice that received no DREADDs-CNO manipulation, or in EC-Tau/hAPP (n=4) mice that were administered a low dose of 0.5mg^-1^/kg^-1^/day^-1^ CNO for 6-weeks. Paired *t*-test: *t* (8) = 1.648, *p* > 0.05. All graphical representations appear as mean ± SEM. Colored bar overlays in Figure 5G-H depict the mean left vs right HIPP 6E10 % values from three immunostained brain sections per mouse. ** *p* < 0.01.

EC-Tau/hAPP mice and hAPP mice that were administered 1mg^-1^/kg^-1^/day^-1^ CNO for 6-weeks exhibited a dramatic reduction in Aβ accumulation within subregions of the hippocampus downstream of the DREADDs-activated right EC (Figure 5E-G). Decreased 6E10+ immunoreactivity was evident throughout the DG, CA3, and CA1 strata (right HIPP: 8.85 ± 1.03 % vs left HIPP: 14.92 ± 1.95 %). Chronic EC DREADDs activation appeared to mainly impact diffuse Aβ accumulation rather than large Aβ deposits, which were present in all HIPP subregions. Hippocampal 6E10+ immunoreactivity was not significantly different between right and left hemispheres in 16-18 month EC-Tau/hAPP (n=2) mice and hAPP (n=4) mice that received no DREADDs-CNO manipulation, or in EC-Tau/hAPP (n=4) mice that were administered a low dose of 0.5mg^-1^/kg^-1^/day^-1^ CNO for 6-weeks (Figure 5H). No significant differences were detected between right and left hippocampal area (mm^2^) sampled for 6E10+ immunostaining comparisons (Supplemental Figure 3 A-D).

These data provide evidence to support the utility of chronic, neuromodulatory intervention in the EC-HIPP network of hAPP/Aβ-expressing mice, as 6E10+ immunoreactivity was significantly reduced in the downstream HIPP after 6-weeks of hM4D_i_ EC DREADDs activation.

### Chronic attenuation of EC neuronal activity *in vivo* reduces tau pathology along the EC-HIPP network

We have previously shown that increased neuronal activity can aggravate tau pathology in EC-Tau mice (32). However, it is unclear whether attenuation of neuronal activity, or attenuation of Aβ-associated hyperactivity in the case of EC-Tau/hAPP mice, can ameliorate local tau pathology or arrest its spread *in vivo*. To address these questions, we performed a series of DAB-IHC staining experiments on horizontal brain sections from chronic hM4D_i_ EC DREADDs activated EC-Tau/hAPP mice (n=4) and EC-Tau (n=5) mice.

We first identified tau aggregates in our tissue by staining for the conformation dependent antibody MC1, which recognizes pathological, abnormally conformed tau (52). Initial MC1+ immunostaining revealed two distinct degrees of pathological tau progression in our mice: Early Tau pathology and Advanced Tau pathology. Mice exhibiting Early Tau pathology have restricted MC1+ immunostaining in parasubiculum (PaS) and EC, with little progression into the hippocampus at 16-months. Mice with Advanced Tau pathology exhibit MC1+ immunostaining along the EC-HIPP network, including progression into the dentate gyrus (DG) and CA1 of the hippocampus (Figure 6). MC1+ immunostaining revealed selective decreases in abnormally conformed tau along the EC-HIPP network in our mice (Figure 6). Representative images from EC-Tau mice (Figure 6A-E) and EC-Tau/hAPP mice (Figure 6F-J) exhibiting Early Tau pathology showed decreased MC1+ immunoreactivity within neuropil and cell bodies of the DREADDs activated rPaS and rEC. These decreases were clearly evident in EC-Tau/hAPP mice (Figure 6G-H), as they possessed a greater number of somatic MC1+ immunostaining in PaS and EC than EC-Tau mice (Figure 6B-C). Notably, MC1+ immunoreactivity was not decreased in the most lateral portion of the rLEC bordering the perirhinal cortex (Figure 6H), an area lacking DREADDs-expression (Figure 6G). A subtle reduction in MC1+ immunostaining was visible in DREADDs-activated rPaS and rEC neuropil (Figure 6B-C). Hemisphere differences in HIPP MC1+ immunostaining could not be detected in mice exhibiting Early Tau pathology, as pathological tau had not significantly progressed into these regions (Figure 6D-E, I-J). Representative MC1+ immunostained sections are shown for EC-Tau mice (Figure 6K-O) and EC-Tau/hAPP mice (Figure 6P-T) exhibiting Advanced Tau pathology. Decreased MC1+ immunostaining could be seen in ipsilateral, downstream DG granule cells, CA1 pyramidal cells and Subiculum in hM4D_i_ EC DREADDs activated EC-Tau mice (Figure 6N-O). Selective decreases in MC1+ immunostaining could also be seen along the EC-HIPP network in EC-Tau/hAPP mice with Advanced Tau pathology, with notable reductions in somatic MC1+ immunostaining in rPaS and rEC (Figure 6Q-R) in the DREADDs activated right hemisphere, as well as in rCA1 pyramidal cells (Figure 6T).

**Figure 6.**
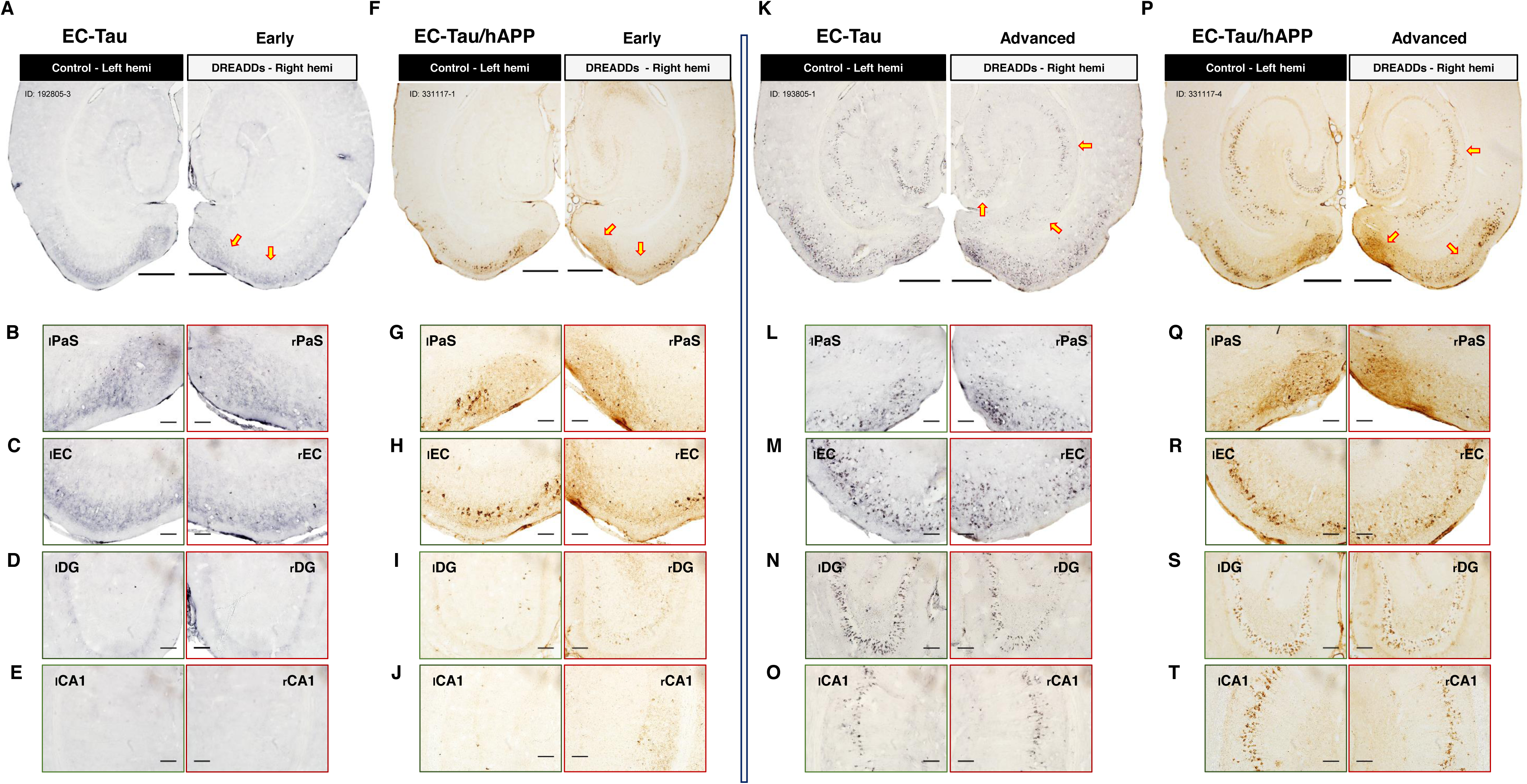
Chronic hM4D_i_ EC DREADDS activation reduces MC1+ staining along the EC-HIPP network. DAB-IHC was performed on horizontal brain sections from 16-month EC-Tau/hAPP (n=4) mice and age-matched, littermate EC-Tau (n=5) mice to evaluate the impact of chronic hM4D_i_ EC DREADDs activation on abnormally conformed tau (MC1). MC1+ immunoreactivity patterns along the EC-HIPP network revealed a distinct separation between mice exhibiting early tau pathology, localized primarily within the EC but not yet in the HIPP, and mice exhibiting significant tau pathology in both the EC and HIPP. We refer to these mice as exhibiting Early Tau Pathology and Advanced Tau Pathology, respectively. ***A.*** EC-Tau, Early Tau pathology; ***F.*** EC-Tau/hAPP, Early Tau pathology; ***K.*** EC-Tau, Advanced Tau pathology; ***P.*** EC-Tau/hAPP, Advanced Tau pathology. Scale bars, 500µm. ***A-E.*** A subtle reduction in MC1+ neuropil staining could be detected in the right PaS and EC of an EC-Tau mouse. ***F-J.*** A clear reduction in MC1+ cell bodies within the right PaS and EC was seen in an EC-Tau/hAPP mouse. ***K-O.*** Reduced MC1+ somatic staining was clearly evident in the DG granule cell layer and CA1 pyramidal cell layer of an EC-Tau mouse with Advanced Tau pathology. ***P-T.*** Reduced MC1+ somatic staining was detected in PaS, EC and CA1 of an EC-Tau/hAPP mouse with Advanced Tau pathology. Yellow arrows indicate areas of reduced MC1+ staining. All images are representative for human tau-expressing mice that fall into Early Tau pathology and Advanced Tau pathology groupings. Magnified images: scale bars, 100µm.

We then processed adjacent tissue sections from experimental mice to identify phosphorylated tau using the phosphotau-specific antibody AT8 (Ser^202^/Thr^205^) (53). Reductions in somatodendritic phosphotau accumulation was also identified along the EC-HIPP network of our hM4D_i_ EC DREADDs activated EC-Tau/hAPP mice and EC-Tau mice (Figure 7). EC-Tau mice exhibiting Early Tau pathology (Figure 7A) showed reduced AT8+ immunoreactivity in the rPaS (Figure 7B) and rEC (Figure 7C) compared to analogous regions in the contralateral hemisphere. EC-Tau/hAPP mice exhibiting Early Tau pathology showed a similar reduction in AT8+ immunostaining (Figure 7F-H). As significant tau pathology had not progressed into the HIPP of these mice at 16-months, no hemisphere differences in HIPP subfield AT8+ immunoreactivity were detected (Figure 7D-E, I-J). EC-Tau mice (Figure 7K) and EC-Tau/hAPP mice (Figure 7P) exhibiting Advanced Tau pathology showed reductions in AT8+ cell bodies and neurites within the rPaS, rEC and rHIPP subfields. Somatic AT8+ immunostaining was reduced in the rSub, rDG granule cells and rCA1 pyramidal cells of EC-Tau mice (Figure 7M-O). EC-Tau/hAPP mice showed reduced AT8+ immunostaining in rPaS and rEC of the DREADDs activated right hemisphere, in addition to downstream CA1 pyramidal cells (Figure 7T). We also stained adjacent tissue sections from experimental mice with the antibody CP27 (human Tau^130-150^) to detect the distribution of total human tau along the EC-HIPP network. Similar to MC1+ and AT8+ immunostaining, CP27+ immunostaining revealed selective decreases in total human tau along the EC-HIPP network in the DREADDs activated right hemisphere of our mice (Supplemental Figure 5).

**Figure 7.**
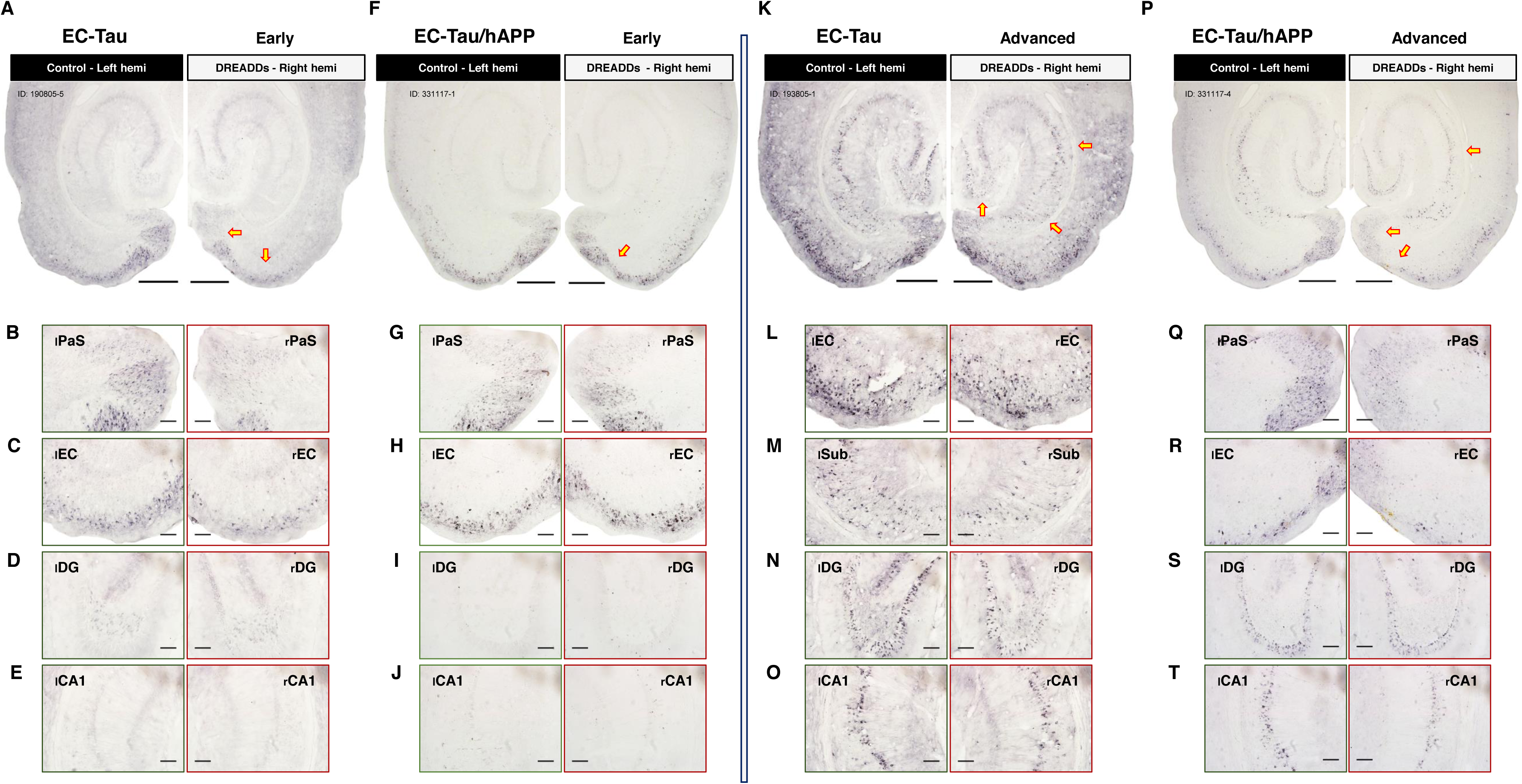
Chronic hM4D_i_ EC DREADDS activation reduces AT8+ staining along the EC-HIPP network. Adjacent tissue sections from hM4D_i_ EC DREADDs activated mice were processed for AT8+ immunostaining experiments to examine the distribution of hyperphosphorylated tau expression along the EC-HIPP network. Horizontal brain sections from 16-month EC-Tau/hAPP (n=4) mice and age-matched, littermate EC-Tau (n=5) mice were processed in parallel. ***A.*** EC-Tau, Early Tau pathology; ***F.*** EC-Tau/hAPP, Early Tau pathology; ***K.*** EC-Tau, Advanced Tau pathology; ***P.*** EC-Tau/hAPP, Advanced Tau pathology. Scale bars, 500µm. ***B-E***. Chronic hM4D_i_ EC DREADDs activation resulted in reduced AT8+ neuronal staining in the right hemisphere PaS and EC in an EC-Tau mouse exhibiting Early Tau pathology. ***F-J***. Reduced AT8+ immunostaining is shown in the rPaS and rEC of an EC-Tau/hAPP mouse exhibiting Early Tau pathology. ***K-O***. Reduced somatic AT8+ immunostaining was evident in the rSub, rDG granule cell layer and rCA1 pyramidal cell layer of an EC-Tau mouse with Advanced Tau pathology. ***P-T.*** Reduced somatic AT8+ immunostaining was evident in the rPaS, rEC and rCA1 pyramidal cell layer of an EC-Tau/hAPP mouse exhibiting Advanced Tau pathology. Yellow arrows indicate areas of reduced AT8+ immunostaining. All images are representative for human tau-expressing mice that fall into Early Tau pathology and Advanced Tau pathology groupings. Magnified images: scale bars, 100µm.

The extent to which chronic hM4D_i_ EC DREADDs activation reduced pathological tau immunoreactivity in the hippocampus varied across mice, with some mice showing strong effects in some regions and others subtle effects. In addition, reduced tau pathology in certain mice was readily detected using some, but not all, tau antibodies. This effect is illustrated in a 16-month DREADDs-activated EC-Tau/hAPP mouse, where reduced tau aggregation is apparent in CP27+ immunostaining, but not in MC1+ or AT8+ immunostaining (Figure 8A). To examine the downstream impact of attenuated EC neuronal activity on hippocampal tau pathology, we performed threshold analysis for each pathological tau marker in the DG, CA1 and SUB. We then calculated the % area of MC1+, AT8+ and CP27+ above threshold for each ROI, similar to methods described in Figure 5G-H analysis. Finally, we generated a right hemisphere versus left hemisphere ratio for each immunostained section and pooled them according to hippocampal subfield (Figure 8B) and pathological tau marker (Figure 8C). Values less than 1 indicate reduced tau accumulation in the treated right hemisphere, while values greater than 1 indicate increased tau accumulation in the right hemisphere. Plotting the right versus left hemisphere ratios by region revealed an effect of hM4D_i_ EC DREADDs activation on tau pathology in the DG (0.8029 ± 0.067) and CA1 (0.7598 ± 0.104), but not the Sub (1.0000 ± 0.117). Plotting the right versus left hemisphere ratios by tau marker revealed the strongest effect of hM4D_i_ EC DREADDs activation on CP27+ immunostaining (0.7714 ± 0.915). AT8+ immunostaining was trending (0.7963 ± 0.097), while MC1+ immunostaining did not show an effect (0.9951 ± 0.108).

**Figure 8.**
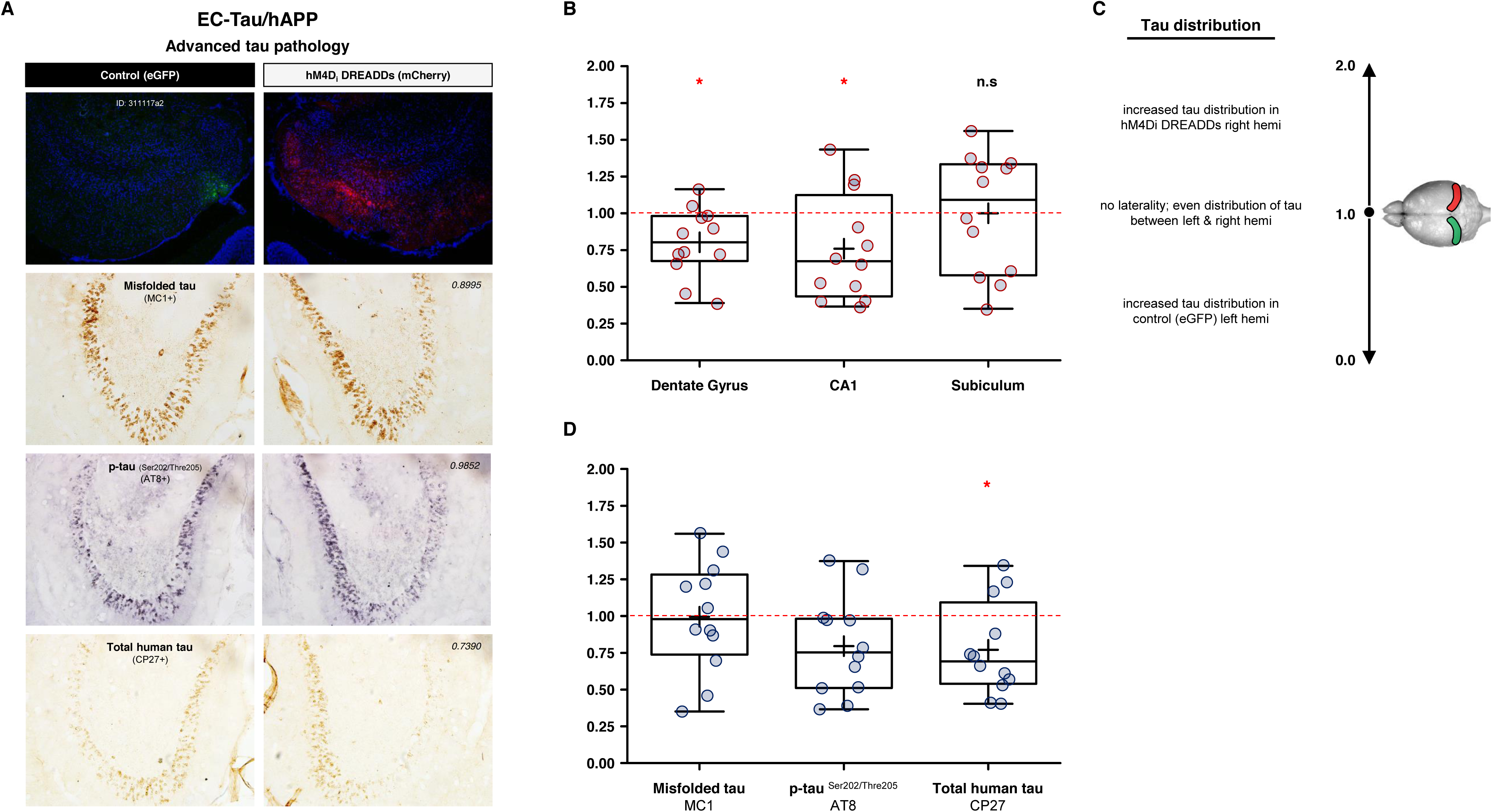
Chronic hM4D_i_ EC DREADDS activation selectively reduces tau pathology in downstream hippocampus. Semi-quantitative analysis of tau pathology expression in HIPP was performed on MC1+, AT8+ and CP27+ immunostained tissue sections of mice exhibiting advanced tau pathology. Within-subject comparisons of tau immunoreactivity (right versus left hemisphere) were calculated per hippocampal region of interest (CA1, DG and Sub) and then plotted as a ratio to reveal any laterality in pathological tau accumulation. Chronic activation of hM4D_i_ EC DREADDs in the right hemisphere was associated with reduced MC1+, AT8+ and CP27+ immunostaining in DG granule cells. DREADDs activation was also associated with reduced AT8+ and CP27+ immunostaining in CA1 pyramidal cells. Box plots depict the mean ± SEM of right versus left hemisphere % Area ratios for experimental mice exhibiting advanced tau pathology (Total, n=4 mice: EC-Tau, n=2; EC-Tau/hAPP, n=2). Black boxes, MC1; Gray, AT8; White, CP27. Transparent, colored overlays depict individual ratio values for each mouse. Hashed line (1.0 ratio) represents equal right versus left hemisphere tau immunoreactivity.

In conclusion, our experimental data strongly support the emerging hypothesis that hAPP/Aβ accumulation in the brain is associated with aberrant neuronal activity and network impairment. Our data describe a role for Aβ-associated neuronal hyperactivity in accelerating tau pathology along a well-characterized neuronal network that is vulnerable to AD pathology and neurodegeneration. We further show that hyperactivity in this network can be targeted via chronic chemogenetic activation to arrest the accumulation of both hAPP/Aβ and pathological tau along the EC-HIPP network *in vivo*.

## Discussion

The progressive, stereotypical spread of pathological tau along neuronal circuits in AD is an active area of intense investigation. However, the underlying mechanisms that potentiate pathological tau spread in AD are currently unresolved. We, and others, have proposed that increased neuronal activity can exacerbate tauopathy either by promoting tau release from neurons or by facilitating its uptake in synaptically connected neurons (32, 37). To model the functional interactions between hAPP/Aβ and hTau in a well-characterized neuronal circuit *in vivo*, we crossed the Aβ-generating hAPP/J20 _(Swedish/Indiana)_ mouse line (21, 54) with the EC-Tau mouse line, wherein mutant hTau _(P301L)_ is predominantly expressed in the EC (10, 11). Network dysfunction and impaired cognitive performance have previously been described in relatively young hAPP mice (21, 22, 48). Mutant hAPP expression in the EC-Tau/hAPP mouse thus provides a strong *in vivo* platform to model the impact of Aβ-associated hyperactivity on tau pathology. At 16 months, somatodendritic MC1+ immunostaining was increased in the EC and DG (Figure 1C-E). By 23-months, the distribution and severity of MC1+ immunostaining resembled that of much older (30+ month) EC-Tau mice (Figure 1F) (39, 55), suggesting that hAPP/Aβ plays a significant role in promoting the intercellular transfer of pathological tau *in vivo*. Our data supports previously published findings in similar mouse models, where robust hAPP/Aβ accumulation was associated with accelerated tauopathy along the EC-HIPP network (36-38).

We hypothesized that increased Aβ production or accumulation may trigger an intermediate, non-pathogenic cascade of events that impact tau. Indeed, substantial evidence exists to support the hypothesis that increased Aβ leads to neuronal hyperactivity and large-scale network dysfunction in the brain (24). This hypothesis may partially explain the accelerated progression of pathological tau in AD and in mouse models of AD pathology. hAPP/Aβ accumulation has been linked to the appearance of epileptiform-like network activity in the brain at a relatively young age (21, 22, 56). This aberrant network activity is associated with changes in inhibitory neuron profiles and remodeling of the DG. More recently, hyperactive neurons have been shown to disproportionately cluster around Aβ plaques in cortex (20), where elevated concentrations of soluble Aβ oligomers may collect (28). Motivated by these observations and our findings (Figure 1), we examined single-unit activity and LFPs in the EC of 16-month EC-Tau/hAPP mice and their age-matched littermates (Figure 2-3). Aβ-generating (EC-Tau/hAPP and hAPP) mice exhibited significant EC single-unit hyperactivity and network dysfunction, characterized by increased average firing rates, decreased median ISIs (Figure 2B-C) and decreased % theta power compared to control mice (Figure 3A-C). Aβ-associated EC hyperactivity was not attributed to oversampling of neuronal bursting in the hAPP-expressing mice, as no significant group differences were found in the % of EC bursting activity recorded (One-way ANOVA: F_(3)_ = 2.426, *p* = 0.1034) (Table 1). Plotting the average firing rates of pooled EC neurons as a function of spike width revealed a distinct population of hyperactive neurons in Aβ-generating mice that was distributed among putative interneurons (NS) and excitatory cells (WS) (Figure 2D-G). This increased proportion of hyperactive cells was not due to oversampling of NS neurons in Aβ-generating mice (NS, n=212) versus Non Aβ-generating mice (NS, n=196) (Table 1). Additionally, Aβ-associated EC hyperactivity and theta impairment was also present after applying a minimum speed filter to the datasets, which removed neuronal activity during bouts of immobility (Supplemental Figure 1). These data support previous findings describing Aβ-associated interneuron dysfunction and network hyperactivity (21, 22, 57), and extend them to putative inhibitory and excitatory neurons in EC. hAPP/Aβ-associated disruptions in theta oscillatory activity are also in line with previous reports (43, 45). Finally, we did not detect hAPP/Aβ-mediated effects on behavioral activity during open field recording sessions (Figure 4). This is in contrast to several reports describing locomotor hyperactivity in the hAPP/J20 mouse line (22, 48, 49). We predict that this may have been due to increased motivational drive to actively explore the arena in our mice, as sucrose pellets were administered during recording sessions to encourage foraging behavior and arena coverage. Repeated handling and acclimation to the recording paradigm may have also reduced innate anxiety-like behavior in our mice, leading to similar behavioral activity patterns. Importantly, we can conclude that Aβ-associated neuronal hyperactivity and network dysfunction are not due to gross locomotor differences in our mice.

Several lines of evidence now support a role for increased neuronal activity in both Aβ accumulation and accelerated tau pathology *in vivo*. Stimulation of the perforant pathway results in increased Aβ concentrations in hippocampal interstitial fluid (ISF) (30), and increased Aβ deposition in downstream DG (outer molecular layer) (33). Chemogenetic stimulation of cortical neurons is also associated with increased deposition of mature amyloid plaques (34). Likewise, stimulating neuronal activity results in increased ISF hTau concentrations (31), and chronic stimulation of EC neurons enhances local tauopathy and accelerates neurodegeneration (32). These data suggest that Aβ-associated hyperactivity can impact pathological tau progression along a defined neural circuit while simultaneously driving increased hAPP/Aβ release. The afflicted circuit could then potentially be recruited into a harmful, positive feedback loop that drives aggressive pathological hAPP/Aβ and tau aggregation in the neuronal network, leading to cognitive impairment and cell death. Aged mice with significant tauopathy along the EC-HIPP network exhibit deficits in spatially modulated grid cell function and impaired spatial learning and memory, as well as excitatory neuron loss, independent of Aβ pathology (39, 55).

Surprisingly, the increased aggregation of pathological tau in EC did not appear to affect measures of hyperactivity or network function in 16-month EC-Tau mice. Single-unit average firing rates, ISI medians and % theta power were remarkably similar in EC-Tau/hAPP and hAPP mice, and in EC-Tau and non-transgenic Control mice. These findings support a previous report from our lab describing normal EC single-unit function in 14-month EC-Tau mice, where average firing rates and SM cell firing patterns matched that of non-transgenic controls (39). These data suggest that the accumulation of tau in this model does not strongly impact EC neuronal activity by 16-months *in vivo*. However, these data should be carefully interpreted, as EC-hTau overexpression has been linked to hypometabolism in ~9-month mice (36) and has recently been shown to blunt Aβ-associated hyperexcitability *in vitro* and *in vivo* (58, 59). It is possible then that detection of hTau-mediated effects on neuronal activity is partly dependent on the sensitivity of the assays used to measure them and degree of pathological severity. Our analysis in 16-month animals revealed blunted hyperactivity in NS neurons of EC-Tau/hAPP mice (Figure 2F), which may represent an early, synergistic effect of Aβ on tau-mediated inhibitory interneuron dysfunction that could precede subsequent impairments in excitatory neurons and gross network function. We have shown that aged (30+ month) EC-Tau mice exhibit significant hypoactivity in excitatory MEC grid cells (39). Thus, we predict that divergent hAPP/Aβ and hTau effects on neuronal activity would be readily observed in the EC-HIPP network of aged EC-Tau/hAPP mice.

We implemented a chemogenetic approach in our studies to combat the aggressive progression of Aβ-associated EC hyperactivity on Aβ and tau pathology *in vivo*. hM4D_i_ DREADDs were targeted to EC principal neurons based on the finding that WS (wide-spiking) cells showed hyperactivity in both hAPP mice and EC-Tau/hAPP mice (Figure 2F-G). Importantly, DREADDs-mediated neuromodulation has been shown to reduce local Aβ deposition in cortex (hM4D_i_) (34) and facilitate the transfer of hTau into distal post-synaptic cell populations (hM3D_q_) (13). Using a within-subjects design, we probed for hAPP/Aβ and pathological tau immunoreactivity in both the ipsilateral, downstream hippocampus and in the contralateral hippocampus. After 6-weeks of hM4D_i_ EC DREADDs activation, we found a marked reduction in 6E10+ immunoreactivity within ipsilateral hippocampus of EC-Tau/hAPP and hAPP mice, supporting previous reports linking stimulated neuronal activity to Aβ release and Aβ pathology (Figure 5E-H) (30, 33, 34, 60, 61). Chronic hM4D_i_ EC DREADDs activation in hTau-expressing mice also led to selective reductions in pathological tau immunoreactivity within ipsilateral, downstream hippocampal subfields, as well as local regions where hM4D_i_ DREADDs were expressed (Figures 6-7, Supplemental Figure 5). Extensive somatodendritic tau aggregates were readily observed within the hippocampus in a subset of mice (characterized as exhibiting ‘Advanced’ tau pathology). To visualize trends in hemispheric tau differences according to hippocampal subfield or tau marker, we pooled right versus left hemisphere ratio values and compared group means to a hypothetical mean of 1.0, which would indicate equal distribution of pathological tau in left and right hemisphere ROIs (Figure 8). We consistently observed reduced tau pathology in ipsilateral rDG granule cells and rCA1 pyramidal neurons, while no group trends were found in rSub (Figure 8B). These data are the first to demonstrate that attenuated neuronal activity can reduce pathological tau *in vivo*, and support previous reports showing that stimulated neuronal activity can increase tau release and tauopathy in AD mouse models (12, 13, 31, 32).

Recent concerns have been raised in the literature regarding the utility of CNO as an inert DREADDs ligand that easily permeates the blood-brain barrier (BBB) (62-64): for review, see (65). In our studies, we confirmed chronic EC DREADDs activation *in vivo* using recording metrics derived from single, high-dose CNO injection studies. Acute hM4D_i_ DREADDs activation resulted in decreased EC neuronal activity beginning at ~20 min, with a maximal response at ~30 min that lasted for at least 4 hr (Supplemental Figure 2A-B). This data is supported by previous *in vivo* research showing strong hM4D_i_ activation in EC 30 min post-CNO, with activity levels recovering towards baseline by 12 hr (66). Acute hM4D_i_ DREADDs activation also led to a robust decrease in % theta power in EC lasting over an hour (Supplemental Figure 2D-E). Therefore, we tracked chronic EC DREADDs activation *in vivo* by analyzing total spike counts and % theta power (Figure 5A-B), rather than measuring levels of CNO or converted clozapine in peripheral blood/plasma. Consistent with our single CNO injection findings, total spike counts and % theta power were reduced over chronic CNO treatment, supporting the utility of long-term CNO delivery in indwelling osmotic minipumps to activate DREADDs *in vivo* (see also (67)).

Expression of hM4D_i_ DREADDs was restricted to the right hemisphere of our mice and did not crossover into the contralateral, left hemisphere (Figure 5C-D and Supplemental Figure 4). Thus, we were able to discriminate the effects of chronic EC DREADDs activation on pathology in ipsilateral, downstream HIPP (right hemisphere) and directly compare it to pathology in the contralateral, control left HIPP. We predict that any off-target effects of chronic, converted CNO-to-clozapine on hAPP/Aβ and tau pathology would have impacted both left and right hemispheres in our experimental mice, as continuous 6-week delivery of 1mg^-1^/kg^-1^/day^-1^ CNO was performed using minipumps. 6E10+ immunostaining revealed strong decreases in hAPP/Aβ in the ipsilateral rHIPP (Figure 5E-G), downstream of the DREADDs activated rEC. We did not detect hemisphere differences in 6E10+ immunoreactivity in age-matched hAPP mice or EC-Tau/hAPP mice sampled from our colony (no DREADDs manipulation), or in hM4D_i_ EC DREADDs expressing EC-Tau/hAPP mice administered a lower CNO dose (0.5mg^-1^/kg^-1^/day^-1^) (Figure 5H and Supplemental Figure 4H). Furthermore, regional decreases in abnormally conformed tau (MC1+), phosphotau accumulation (AT8+) and total human tau (CP27+) were detected in ipsilateral, rHIPP subfields after chronic hM4D_i_ EC DREADDs activation. The degree to which pathological tau and Aβ accumulation were reduced in downstream HIPP, along with our electrophysiology data showing reduced neuronal activity, is consistent with the hypothesis that ~50-75% maximal DREADDs activation could be achieved with 1mg^-1^/kg^-1^/day^-1^ CNO (68).

Attenuating neuronal hyperactivity and network dysfunction may prove to be a powerful tool in combating impaired cognition in human AD, especially when paired with therapies aimed at alleviating the aggregation and deposition of Aβ and tau. On its own, Aβ-targeted immunotherapy has proven unsuccessful at relieving AD cognitive symptoms, which may be due to ineffective amelioration or the exacerbation of neuronal hyperactivity (29, 69). Indeed, reducing AD pathology-associated neuronal dysfunction with the antiepileptic drug leviteracetam has shown promise in preclinical mouse models of hAPP overexpression (22, 70), and is being tested for efficacy in AD clinical trials. Our data shows that alleviating Aβ-associated EC hyperactivity reduces downstream accumulation of both Aβ and tau pathology in the hippocampus. An important step forward will be to replicate these findings in non-overexpressing hAPP mice (APP ^NL-F/NL-F^), which show early signs of neuronal hyperexcitability *in vitro* and impaired gamma oscillations *in vivo* (71-73). In addition, future studies will be required to determine if relief from pathological Aβ and tau using approaches such as chemogenetics will improve cognitive behavior. Interestingly, reducing murine tau levels has been shown to alleviate locomotor hyperactivity in young hAPP/J20 mice and can blunt pharmacologically induced aberrant over-excitation (49, 74). This would indicate that several potential mechanisms exist in the brain to contribute to impaired neuronal activity in AD, and provide ample avenues for investigation into the etiology of AD progression.

## Author Contributions

These experiments were designed by G.A.R., K.E.D. & S.A.H. All experiments were performed by G.A.R. Data analysis was performed by G.A.R., G.M.B. & S.A.H. The manuscript was written by G.A.R., K.E.D. & S.A.H.

### Acknowledgments

We thank Chukwuma Onyebuenyi, B.S., and Paula Choconta, B.S., for expert technical assistance with electrophysiological data analysis and image processing. We thank Helen Y Figueroa, B.S., for maintaining the EC-Tau/hAPP mouse colonies. We thank Dr. Peter Davies for generously providing tau antibodies. The authors would also like to thank Drs. Wai Huang Yu, Catherine Clelland, Natura Myekua and Tal Nuriel for helpful discussions and comments regarding the manuscript. This work was supported by research grants from the NIH (R01AG050425 to S.A.H. and R01AG050425-Supplement to S.A.H./G.A.R.) and the Alzheimer’s Association (AARFD-17-504409 to G.A.R.).

## Materials and Methods

### Experimental animals

Three transgenic mouse lines were used in these studies to model hallmark AD pathologies in the brain. 1^st^ Line: EC-Tau mice that overexpress human mutant tau (4R0N P301L) predominantly in the EC (10, 11, 75) on a C57BL/6 background. 2^nd^ Line: hAPP/J20 mice that overexpress hAPP with two familial AD mutations (KM670/671NL, Swedish) (V717F, Indiana) (54, 76) on a FVB/N background. 3^rd^ Line: EC-Tau/hAPP mice, which generate both hAPP/Aβ and tau pathology in the brain, were created by crossing the EC-Tau and hAPP mouse lines.

A total of 58 mice, both males and females, were used as experimental animals in these studies. Total mouse numbers per genotype were as follows: non-transgenic controls (Control, n=8), and age-matched, transgenic littermates (EC-Tau, n=6; hAPP, n=9; EC-Tau/hAPP, n=8). Additionally, two mice from each genotype were sampled from our colony at 10-months, 16-months, and 23-months of age for initial pathology studies (n=18) in Figure 1. All mice were housed in a temperature and humidity controlled vivarium at Columbia University Medical Center and maintained on a standard 12 hr light/dark cycle with food and water provided *ad libitum*. All animal experiments were performed during the light phase in accordance with national guidelines (National Institutes of Health) and approved by the Institutional Animal Care and Use Committee of Columbia University.

### Microdrive construction

Microdrives were constructed as described previously (39, 77). Briefly, custom-made reusable 16-channel or 32-channel microdrives (Axona, UK) were outfitted with four to eight tetrodes consisting of twisted, 25 mm thick platinum-iridium wires (California Wires, USA) funneled through a 23 ga stainless steel inner cannula. A 19 ga protective, stainless steel outer cannula was slipped over the inner cannula and secured to the microdrive body via modeling clay. Individual electrodes were wrapped tightly around the exposed wires of the microdrive and coated with a layer of Pelco conductive silver paint (Ted Pella, Inc., USA) prior to sealing of the microdrive body with liquid electrical tape (Gardner Bender, USA). Several hours prior to surgery, the tetrodes were cut to an appropriate length and electroplated with a platinum/gold solution until the impedances dropped within a range of 150-200 ohms.

### DREADDs virus microinjection and tetrode implantation

A total of 30 mice were implanted and recorded for *in vivo* electrophysiology studies (Control, n=7; EC-Tau, n=6; hAPP, n=9; EC-Tau/hAPP, n=8). On the day of electrode implantation, mice were anesthetized with isoflurane (3-4% for induction; 0.5-3% for maintenance) using a multi-channel VetFlo Traditional Anesthesia vaporizer (Kent Scientific) and fixed within a stereotaxic frame (Kopf Instruments). As described previously (39), an incision was made to expose the skull and 3 jeweler’s screws were inserted into the skull to support the microdrive implant. A 2 mm hole was made on each side of the skull at position 3.0-3.1 mm lateral to lambda and ~ 0.2 mm in front of the transverse sinus. Viral delivery of the G_i_-coupled hM4D_i_ DREADD (AAV5-CamKII_α_-hM4D_i_-mCherry, 4.1×10^12^) (Cat No. 50477-AAV5; Addgene, USA) was administered into the right EC via a 33 ga NeuroSyringe (Hamilton, USA) tilted at an angle of 6-7° in the sagittal plane. A control virus (AAV5-CamKII_α_-eGFP, 4.1×10^12^) (Cat No. 50469-AAV5; Addgene, USA) was administered into the contralateral left EC using identical parameters. At this point, an additional screw connected with wire was inserted into the skull, serving as a ground/reference for local field potential (LFP) recordings. The prepared 16-channel microdrive was then tilted at 6-7° on a stereotaxic arm and the tetrodes lowered to 1.0 mm from the surface of the brain (below dura) into the DREADDs-delivered right hemisphere. The microdrive ground wire was then soldered to the skull screw wire and the microdrive was secured with dental cement. Mice were then allowed to recover in a cleaned cage atop a warm heating pad until awake (~45 min) before being transported to the housing facility. Mice received Carprofen (5mg/kg) prior to surgery and post-operatively to reduce pain, in additional to a sterile saline injection (s.c.) to aid in hydration. Recording experiments began approximately one week from the time of surgery.

An additional cohort of 16-month hAPP (n=3) mice were microinjected with virus and implanted with osmotic minipumps for long-term DREADDs activation, but did not receive microdrive implants for electrophysiological recordings.

### In vivo recording: single-unit and local field potential analyses

All mice outfitted with microdrives underwent four to six recording sessions in an arena (70 cm x 70 cm), with one recording session performed per day. Tetrodes for each mouse were moved down 100 µm from their previous position 24 hr prior to the next recording session, allowing stable electrode positioning and a robust sampling of EC neuronal activity for each mouse. Additionally, the arena and visual cue were rotated between sessions.

Neuronal signals from our mice were recorded using the Axona DacqUSB system, and described previously (39). Briefly, recording signals were amplified 10,000 to 30,000 times and bandpass filtered between 0.8 and 6.7 kHz. The LFP was recorded from four channels of each microdrive, amplified 8,000 to 12,000 times, lowpass filtered at 125 Hz and sampled at 250 Hz. 60 Hz noise was eliminated using a Notch filter. Spike sorting was performed offline using TINT cluster-cutting software and Klustakwik automated clustering tool. The resulting clusters were further refined manually and were validated using autocorrelation and cross-correlation functions as additional separation tools. Only cells that produced a minimum of 100 spikes with less than 1% refractory period violations (refractory periods < 1ms) were used for subsequent analysis (41). Single-units with no undershoot in their waveform were discarded (defined as Area Under the Peak < 1.0 µV^2^). Putative excitatory neurons (WS, wide-spiking) were distinguished from putative interneurons (NS, narrow-spiking) by first examining the frequency distribution histogram of pooled EC single-unit spike widths, and then bisecting the waveform spike widths according to the following calculation:

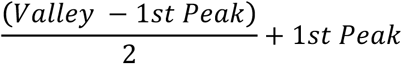

This resulted in a delineation cutoff at 300µs (39, 41). Quantitative measurements of cluster quality were then performed, yielding isolation distance values in Mahalanobis space (78). There were no significant differences between clusters in our experimental groups (Median isolation distances: Control, 7.875; EC-Tau, 8.220; hAPP, 8.670; EC-Tau/hAPP, 8.120: Kruskal-Wallis test, H=7.00, *p* > 0.05).

A total of 1,260 EC neurons were recorded across 20 mice in this study (Control, n=334 single-units (n=7 mice); EC-Tau, n=239 (n=4 mice); hAPP, n=356 (n=5 mice); EC-Tau/hAPP, n=331 (n=4 mice). A breakdown of the analyzed data can be found in Supplemental Table 1.

### Osmotic minipump preparation and surgical implantation

Following EC recording sessions, a subset of EC-Tau/hAPP (n=4) mice and EC-Tau (n=5) mice underwent a second surgical procedure to chronically activate hM4D_i_ EC DREADDs. In addition, a subset of hAPP (n=3) mice that did not receive microdrive implants was subject to chronic hM4D_i_ EC DREADDs activation. Clozapine-n-oxide (CNO; Sigma Aldrich, USA) was dissolved in sterile saline with 0.05%

24 hr prior to surgery, osmotic minipumps (Model 2006) (Alzet, USA) were primed with clozapine-n-oxide (CNO) at a concentration range of 7-10µg/µL in a sterile fume hood (Alzet drug concentration calculator: flow rate, 0.15µL/hr; target dose, 1mg^-1^/kg^-1^/day^-1^). CNO-filled pumps were stored in warm (37°C) sterile saline in a conical tube until surgery.

Mice were once again anesthetized with isoflurane as previously described and the fur clipped at the abdomen. A single injection of marcaine (2mg/kg, 0.05mL) was delivered intradermally into the site of incision ~ 5 min prior to surgery, then a midline incision was made in the abdominal wall and a sterile, CNO-filled minipump was maneuvered into the intraperitoneal (i.p.) cavity. The incision was then closed using an absorbable suture (Henry Schein, USA) for the abdominal layer, followed by closure of the skin with a nylon synthetic non-absorbable suture (Henry Schein, USA). A topical antibiotic was then administered at the surgical site and the mice were allowed to recover in a cleaned cage atop a warm heating pad until awake (~15 min). Mice implanted with minipumps were administered Carprofen (5mg/kg) prior to surgery and post-operatively to help reduce pain. Non-absorbable sutures were removed ~10 days after surgery.

### DAB immunohistochemistry and immunofluorescence imaging

At 6 weeks, all mice were deeply anesthetized with a cocktail of ketamine/xylazine before being transcardially perfused with ice-cold 100mM phosphate-buffered saline (PBS) pH 7.4., followed by 10% formalin (Fisher Scientific, USA). The last recording position for each microdrive-implanted mouse was recorded, and then the microdrive removed. Brains were then harvested and left in 10% formalin overnight, then incubated in 30% sucrose until the brains sank to the bottom of the conical tube (all at 4°C). Horizontal brain sections were sliced (30µm) using a Leica CM3050 S cryostat and stored in cryoprotectant at −20°C until immunostaining procedures. For both immunoperoxidase staining and immunofluorescence imaging, EC-Tau/hAPP brain sections were processed in parallel with sections from EC-Tau and hAPP mice, and non-transgenic Control mice where appropriate. Finally, all tissue sections included for semiquantitative analysis of hAPP/Aβ (Figure 4) and tau (Figure 7) were verified to be within the range of DREADDs mCherry expression by first checking for native fluorescence signal in free-floating sections on an inverted Olympus epifluorescence microscope.

Immunoperoxidase staining was performed using a Mouse-on-Mouse kit (Vector Laboratories, USA) and modified from previous methodology (11, 39, 55). Briefly, cryoprotectant was washed from the free-floating tissue sections with PBS before quenching endogenous peroxidases with 3% H_2_O_2_. Sections were then blocked with mouse IgG blocking reagent for 1.5 hr at room temperature, followed by overnight incubation at 4°C with either anti-beta amyloid 6E10 (mouse, 1:1,000 dilution of 1mg/mL stock) (Biolegend, USA), anti-tau MC1 (mouse, 1:500; courtesy of Peter Davies, Albert Einstein College of Medicine), biotinylated anti-phospho tau^Ser202/Thr205^ AT8 (mouse, 1:500) (Thermo Scientific, USA) or anti-human tau CP27 (mouse, 1:500; courtesy of Peter Davies, Albert Einstein College of Medicine). Tissue sections were then rinsed in PBS and incubated for ~20 min at room temperature in a working solution of biotinylated anti-mouse IgG reagent (excluding the biotinylated AT8-labeled sections). After several PBS rinses, sections were then incubated in an avidin-biotin conjugate for 10 min before being developed in H_2_O containing 3’3’-diaminobenzidine (DAB) hydrochloride and urea hydrogen peroxide (Sigma Aldrich, USA). After staining was completed, tissue sections were mounted onto glass Superfrost Plus slides (Fisher Scientific, USA), allowed to air dry completely, and then dehydrated in ethanol and cleared with xylenes before being coverslipped.

Horizontal tissue sections used to visualize native DREADDs mCherry and eGFP expression in the brain were first rinsed in PBS containing 0.3%Triton X-100 (Sigma Aldrich, USA) (PBST) and then incubated in a working solution of Hoechst 33342 dye (5µg/mL) (Thermo Scientific, USA) to stain cell nuclei for 10 min at room temperature. Subsequent washes in PBST were followed by mounting the tissue onto Superfrost Plus slides and coverslipping using SlowFade gold anti-fade reagent (Life Technologies, USA). All slides were stored in the dark at 4°C until imaging.

### DAB IHC image analysis

Immunoperoxidase-stained tissue sections were analyzed under bright field microscopy using an Olympus BX53 upright microscope. Digital images were acquired using an Olympus DP72 12.5 Megapixel cooled digital color camera connected to a Dell computer running the Olympus cellSens software platform. Image files were then coded and analyzed by an investigator blinded to genotype and saved to a Dell Optiplex 7020 (79).

Semiquantitative analysis of MC1+ cell counts in EC and DG was performed in horizontal brain sections from 16-month EC-Tau/hAPP mice (n=4) and EC-Tau mice (n=5) (Figure 1E). MC1+ cells within defined ROIs (EC and DG granule cell layers) were counted using the multi-point tool and saved via the ROI Manager. The estimated total number of MC1+ cells/mm^2^ was then calculated per left and right hemisphere ROI in each section and averaged across three sections per mouse to generate a representative value. Finally, MC1+ cells/mm^2^ values for each ROI were pooled and compared across genotype. Only cells with clear somatodendritic accumulation of MC1+ immunoreactivity were included in our analysis.

For DAB IHC threshold analysis, minimum threshold values for 8-bit, grayscale images of immunoperoxidase-stained sections were adjusted interactively under blinded conditions in Fiji using an over/under display mode (red represents pixels above threshold) (Supplemental Figure 3A & 3F). A region of interest (ROI) was then defined within each image and saved via the ROI Manager. The total immunoreactivity for each antibody marker above threshold was saved as a percentage of total ROI area (mm^2^) and used to compare pathological accumulation of hAPP/Aβ and tau in the right versus left hemispheres. The same minimum threshold value was applied for each pair of images (right vs left ROIs) used in our analysis. For all DAB IHC experiments, three immunostained tissue sections were analyzed per mouse and averaged to generate a representative value reflecting hAPP/Aβ and tau pathology.

### Behavioral analysis

Animal performance during *in vivo* recording sessions was assessed by tracking the position of an infrared LED on the head stage (position sampling frequency, 50 Hz) by means of an overhead video camera. The position data were first centered around the point (0, 0), and then speed filtered where noted, with only speeds of 5 cm/sec or more included in the analysis. Tracking artifacts were removed by deleting samples greater than 100 cm/sec and missing positions were interpolated with total durations less than 1 sec, and then smoothing the path using a moving average filter with a window size of 300 ms (15 samples on each side). The total distance traveled in the arena (m), % of arena coverage, and average speed (cm/sec) during exploration served as dependent measures of interest. For % of arena coverage, the processed position data was plotted and converted to black and white. Using the ‘bwarea’ function in MATLAB, we calculated the total area traveled by the mouse using the binary image, and then divided this area by the total area of the arena. For each dependent measure, an average value per mouse was generated by collapsing three individual recording sessions that occurred over three days. Values for the entire session were then pooled per genotype and compared, or split into the following time bins and compared: the first 5 min, 10 min and 15 min of the total recording session (30 min).

### Tools for semi-automated data analysis

#### BatchTINTV3

BatchTINTV3 is a graphical user interface (GUI) written in Python, and created as an end-user friendly batch processing solution to complement Axona’s command line interface. BatchTINTV3 sorts the spike data of each session in a chosen directory using KlustaKwik. The BatchTINTV3 code is freely available and hosted in the following GitHub repository: https://github.com/hussainilab/BatchTINTV3.

#### BatchPowerSpectrum

The percent power values were calculated in MATLAB. A Welch’s power spectrum density (PSD) estimate of the LFP was calculated via the ‘pwelch’ function. Using the PSD, the average powers in each of the desired frequency bands were calculated with the ‘bandpower’ function. The average power of each band was then divided by the total power of the signal to produce the percentage power in each of the bands. In situations where the data was speed filtered (5-30 cm/sec), the speed of the animal was calculated and then interpolated so there was a speed value for each LFP value. Then the LFP values were chunked to contain consecutive data points where the mouse’s movements satisfied the minimum and maximum speed requirements. Chunks containing less than 1 second of data were discarded. The aforementioned power calculations were then performed on each of the LFP chunks, and an average of these chunks would yield the final percentage power values for each of the frequency ranges. The BatchPowerSpectrum code is freely available and hosted in the following GitHub repository: https://github.com/hussainilab/BatchPowerSpectrum.

#### hfoGUI for time-frequency analysis

A time-frequency representation (TFR) of the LFP was visualized using a GUI written in Python called ‘hfoGUI.py’. The GUI allows for complete control of signal filtering and is equipped with various filter types (butterworth, chebyshev type 1, chebyshev type 2, etc.), along with the ability to specify filter order, cutoffs, etc. We used 3rd order butterworth filters in order to maintain consistency with the filter types and order implemented by the Axona data acquisition software/hardware. Specific time windows of the LFP data were selected and subjected to a Stockwell-Transform (s-transform) to visually represent EC-Tau/hAPP and Control genotypes (Figure 3A-B). The hfoGUI code is freely available and hosted in the following GitHub repository: https://github.com/hussainilab/hfoGUI.

### Statistical Analysis

Statistical analyses were performed in GraphPad Prism 7 and Matlab. All datasets were tested for normality using the Shapiro-Wilk test. Datasets where values were not well modeled by a normal distribution were subjected to non-parametric statistical analyses. Unpaired *t-*tests with Welsh’s correction were used to perform semiquantitative comparisons of tau pathology in Figure 1. Two sample Kolmogorov-Smirnov tests were used to compare distributions of average firing rates and Median ISIs in Figure 2. The Kruskal-Wallis *H* test was used to compare the medians of the AVG FRs and Median ISIs across genotypes in Figure 2. One-way ANOVAs were used to compare the LFP group means in Figure 3 and locomotor activity in the open field in Figure 4. A repeated measures ANOVA was used to calculate the difference in % theta power over time in Figure 5 (compared to baseline). Paired *t*-tests were used to compare 6E10+ immunostaining in the DREADDs activated right hemisphere versus Control left hemisphere in Figure 5. One-sample *t*-tests were used to compare averaged right versus left hemisphere ratios for tau markers to a theoretical mean (1.0) in Figure 8. *Post hoc* analyses were performed using Dunnett’s multiple comparison test where noted.

## Supplemental Figure Legends

**Supplemental Figure 1.**
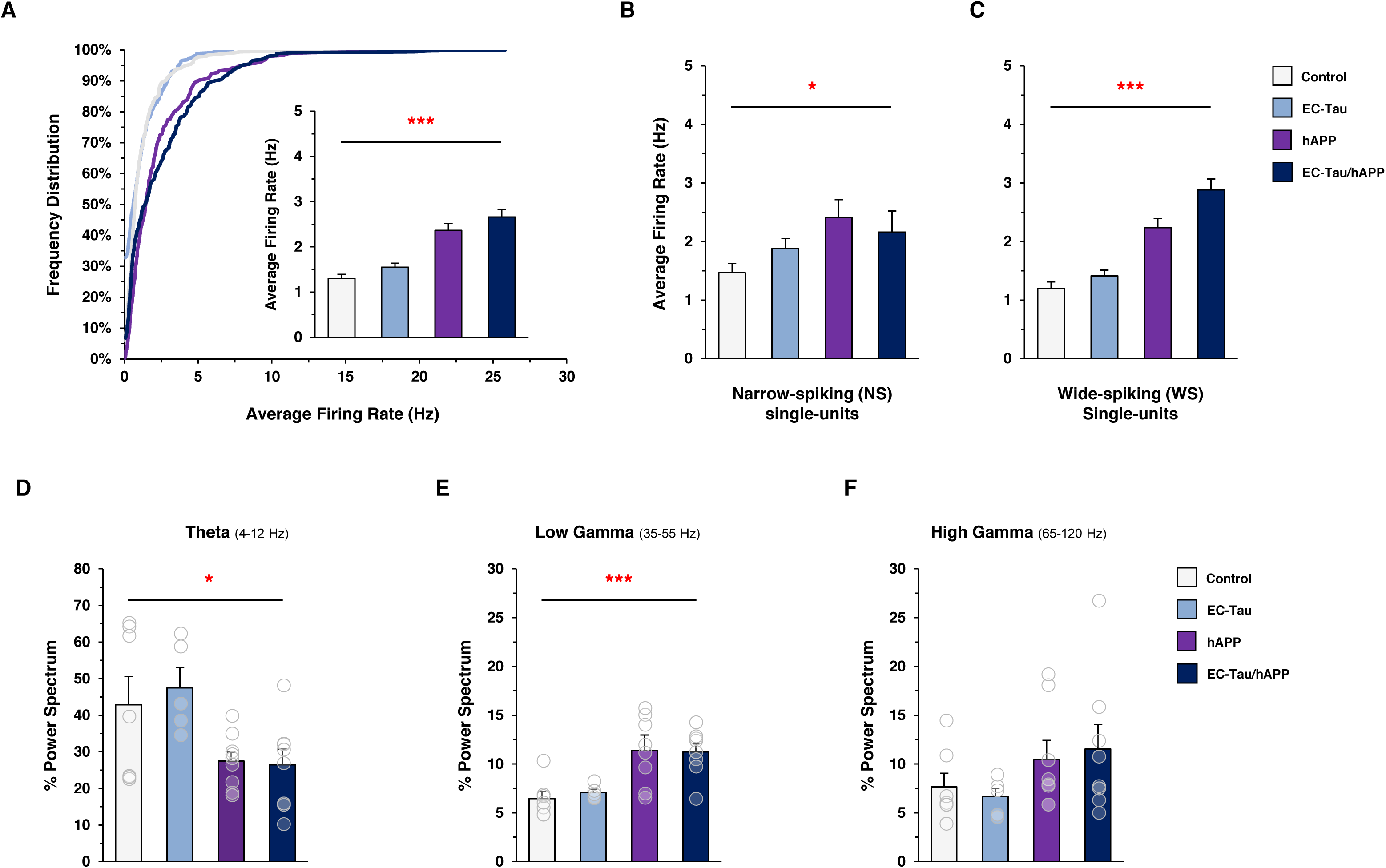
Speed filtered EC single-unit firing rates and network activity. A minimal speed filter (<5cm/sec) was applied to the electrophysiology datasets *post hoc* to remove single-unit spikes and LFP activity that occurred during bouts of immobility. ***A***. The cumulative frequency distributions of pooled, speed-filtered EC single-unit AVG FRs for each genotype are shown. hAPP (n=356 cells; n=5 mice) and EC-Tau/hAPP (n=331 cells; n=4 mice) EC single-units exhibited a clear separation from EC-Tau (n = 239 cells; n=4 mice) and Control (n = 334 cells; n=7 mice) single-units. *Insert*, the speed-filtered, AVG FRs of hAPP EC singe-units were significantly increased compared to Control. Kruskal-Wallis test: H = 61.23, *p* < 0.0001; Dunn’s multiple comparison tests, *p* <0.001. ***B-C***. EC single-units were classified as narrow-spiking (NS) or wide-spiking (WS) cells, as described in Methods and Results. Significant differences were found in the speed-filtered, AVG FRs of NS single-units and WS single-units. NS single-units, H=10.58, *p* < 0.01. WS single-units, H=82.56, *p* < 0.001. ***D***. The averaged, speed-filtered % theta power values of hAPP (27.49 ± 2.34 %) mice and EC-Tau/hAPP (26.44 ± 4.32 %) mice were decreased compared to Control (42.83 ± 7.72 %) mice. One-way ANOVA: F = 3.777, *p* < 0.05. ***E***. Speed-filtered % low gamma power was increased in hAPP (11.38 ± 1.24 %) mice and EC-Tau/hAPP (11.24 ± 0.85 %) mice compared to Control (6.44 ± 0.72 %) mice. One-way ANOVA: F = 7.810, *p* < 0.001. ***F***. No differences were detected across genotype in speed-filtered % high gamma power. Data represented as mean ± SEM. Transparent circles overlaid onto bar graphs depict individual speed-filtered, averaged % power values for each mouse, respectively. * *p* < 0.05; ** *p* < 0.01; *** *p* < 0.001

**Supplemental Figure 2.**
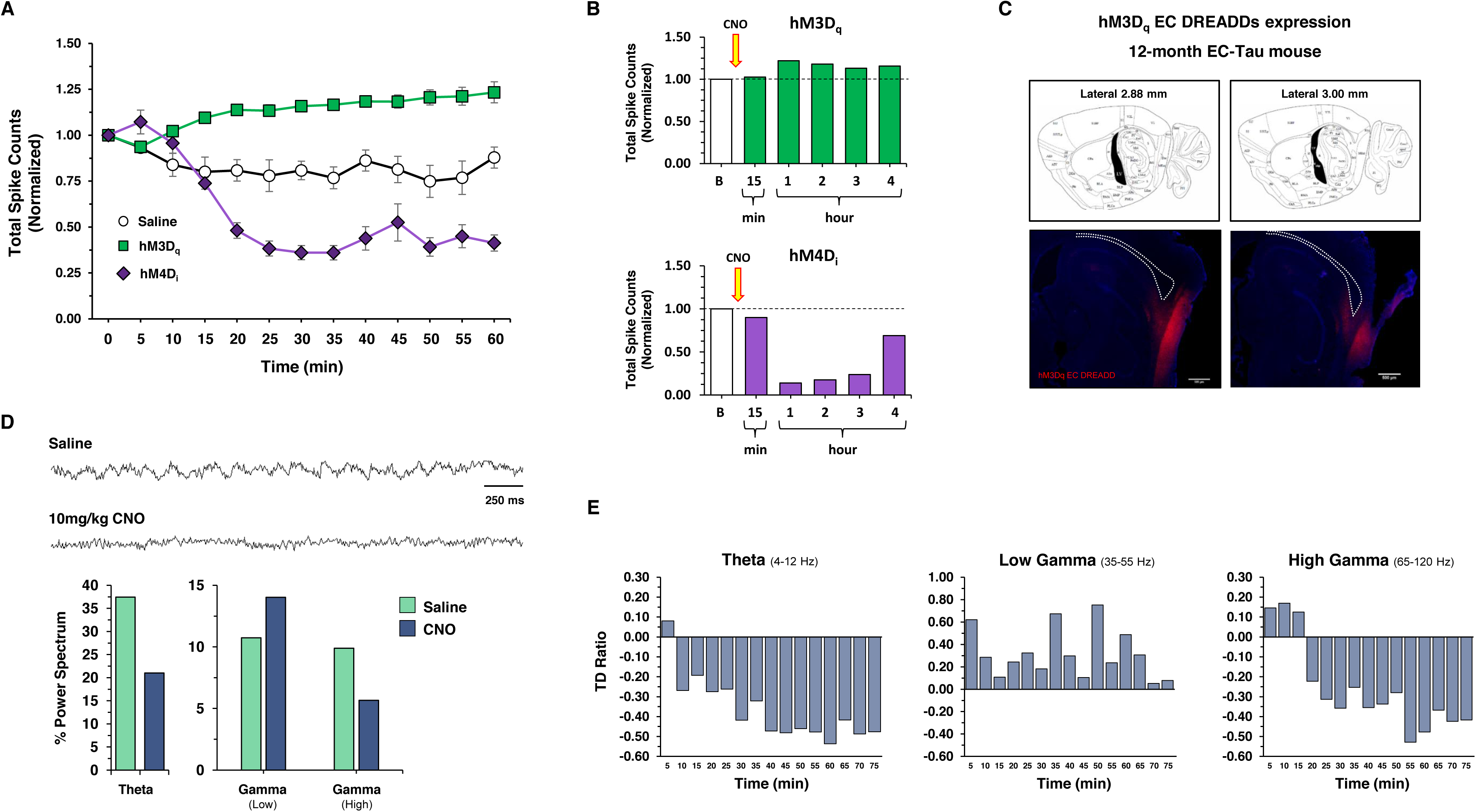
Acute EC DREADDs activation in AD mouse models. Single injections of CNO (i.p.) were used to determine salient recording measures of EC DREADDs activation *in vivo*. ***A-B***. Long-term recordings were first performed in naïve 12-13 month EC-Tau mice to determine the onset and duration of altered single-unit activity due to EC DREADDs activation (hM3D_q_ EC, n=1 mouse; hM4D_i_ EC, n=1 mouse). *Left*, Total spike counts (normalized to baseline measures) are shown for each mouse following CNO and Saline administration (60 min post-treatment). EC DREADDs activation occurs ~20 min post-CNO compared to Saline and lasts at least 4 hr (*right*). Square (green), hM3D_q_ EC DREADDs; Diamond (purple), hM4D_i_ EC DREADDs; Circle (white), Saline. ***C***. Sagittal sections from an EC-Tau mouse expressing hM3D_q_ EC DREADDs is shown. DREADDs-mCherry expression is present along the dorsoventral axis of the EC. Tetrode tract is shown. Scale bars, 500µm. ***D***. The effects of acute hM4D_i_ EC DREADDs activation on LFP measures in EC are shown in a 12-month hAPP/J20 mouse. LFP traces are shown for both CNO and Saline conditions. Notable differences in EC theta modulation were present after 10mg/kg CNO treatment. ***E***. To determine the long-term effects of CNO treatment on LFP measures, the magnitude difference in % theta, % low gamma and % high gamma power versus Saline were calculated. % theta power and % high gamma power in EC were robustly affected by CNO treatment for over an hour. TD Ratio is the normalized Treatment Difference ratio showing magnitude difference of CNO treatment vs Saline.

**Supplemental Figure 3.**
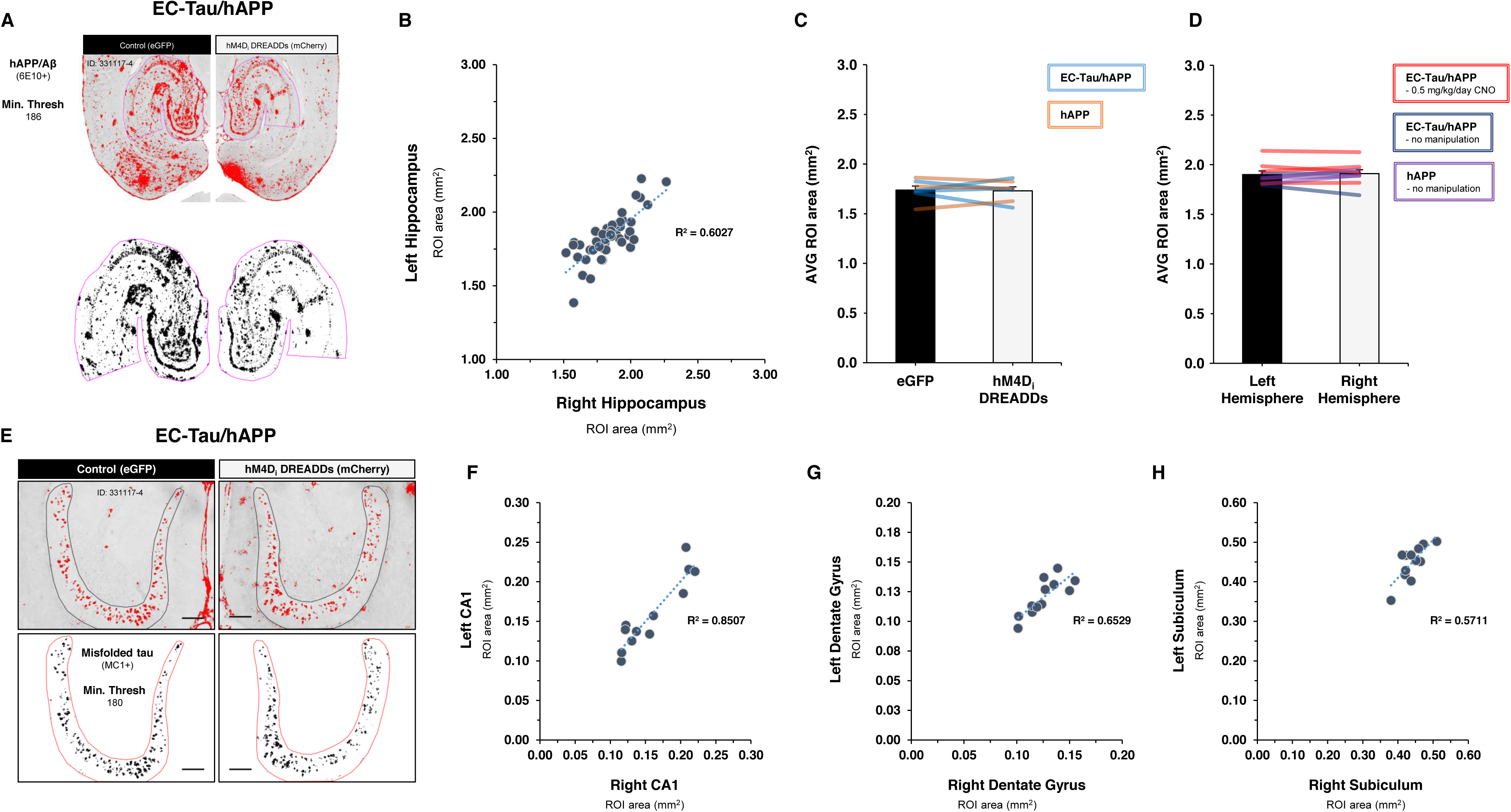
Right vs left hemisphere ROI measures for DAB IHC analysis. The ipsilateral HIPP regions of interest (ROI) area (mm^2^) downstream from the hM4D_i_ DREADDs-expressing EC were compared to contralateral HIPP ROIs. Positive immunoreactivity for each marker was also analyzed within each defined ROI and compared across hemispheres as described in Methods. ***A***. *Top*, 8-bit gray scale images of 6E10+ immunoreactivity in a horizontal brain section from a 16-month EC-Tau/hAPP mouse. A minimum threshold value was first applied to each image (186), and then the ROIs (lHIPP and rHIPP) were defined (magenta). Finally, the % area of 6E10+ immunoreactivity above threshold was used to quantify hAPP/Aβ accumulation in the right and left HIPP ROIs. Scale bars, 500µm. *Bottom*, Higher magnification of 6E10+ immunoreactivity within ROIs. Black pixels depict 6E10+ immunoreactivity above threshold on a white background. Scale bars, 250µm. ***B***. Right vs left hippocampal ROI values (area, mm^2^) for each tissue section analyzed in Figure 4G-H. The coefficient of determination (R^2^, 0.6027) is shown. ***C***. The averaged ROI area values did not differ between the right hemispheres (1.731 ± 0.039 mm^2^) and left hemispheres (1.737 ± 0.039 mm^2^). Paired *t* test: *t* (6) = 0.1557, *p* > 0.05. Three sections averaged per mouse. EC-Tau/hAPP, n=4 mice; hAPP, n=3 mice. ***D***. The averaged ROI area values did not differ between the right hemispheres (1.910 ± 0.040 mm^2^) and the left hemispheres (1.899 ± 0.037 mm^2^) in control conditions. Paired *t* test: *t* (6) = 0.5382, *p* > 0.05. ***E***. 8-bit gray scale images of MC1+ immunoreactivity in the left and right dentate gyrus (DG) of a 16-month EC-Tau/hAPP mouse. A minimum threshold value (180) was applied and the granule cell layers were delineated in black. Within the ROIs, the % area of MC1+ immunoreactivity above threshold was quantified. Accompanying black and white over/under images are shown to emphasis MC1+ signal in DG ROIs. Scale bars, 100µm. ***F-H***. The right versus left ROI values (mm^2^) for hippocampal subregions analyzed were plotted. CA1: R^2^=0.8507; DG granule cells: R^2^=0.6529; Subiculum: R^2^=0.5711. Bar graphs represent mean ± SEM. Colored bar overlays depict the mean right versus left HIPP ROI area (mm^2^) values for each mouse.

**Supplemental Figure 4.**
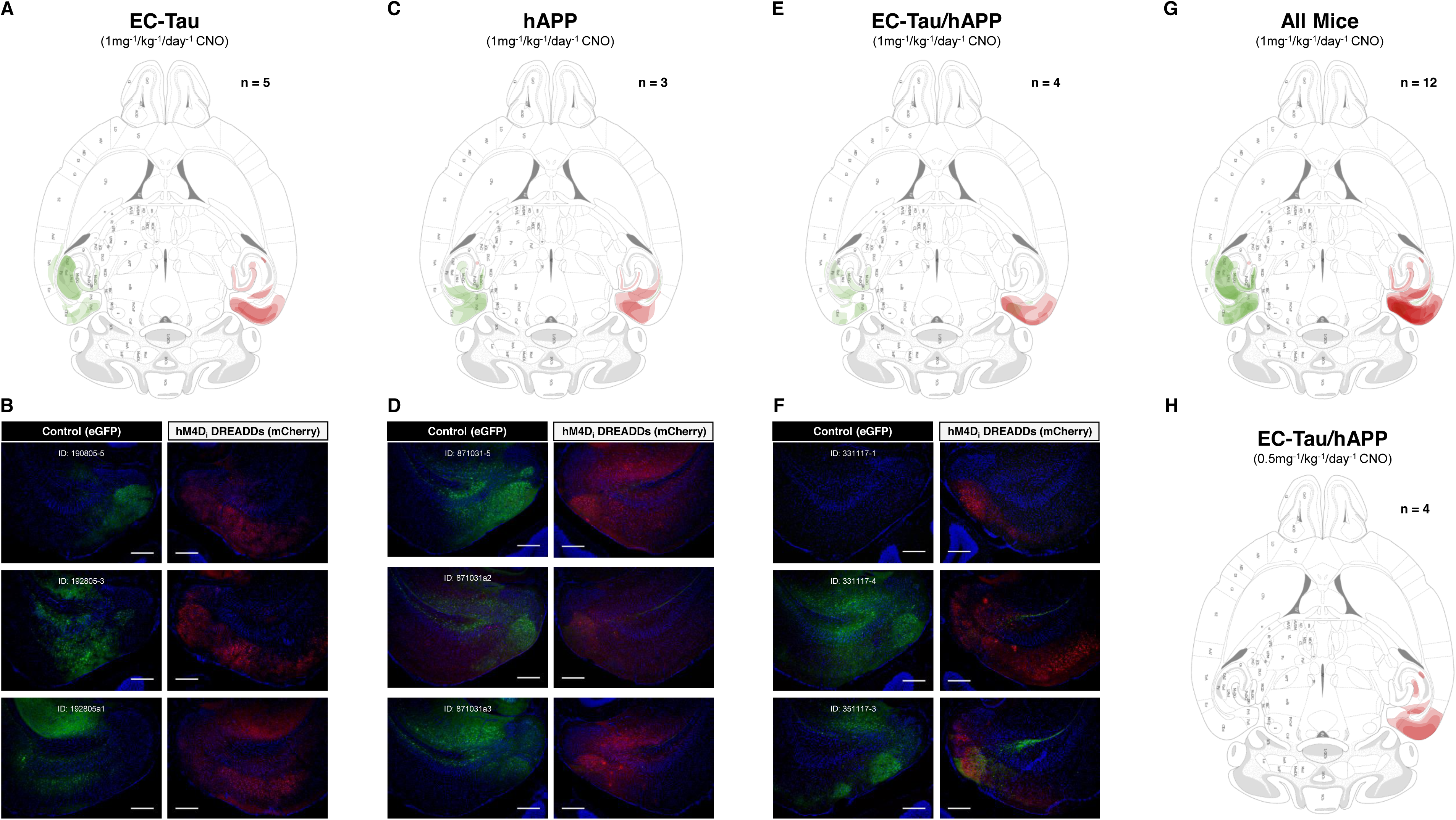
hM4D_i_ EC DREADDs expression in experimental mice. Regional mCherry (hM4D_i_ DREADDS) and eGFP (Control) expression patterns were verified in all mice. For each group, illustrated mCherry and eGFP expression patterns were overlaid onto a mouse stereotaxic brain atlas. ***A***. EC-Tau mice (n=5) exhibit robust hM4D_i_ DREADDs expression throughout the rEC, rPaS and rPrS. mCherry signal was present in the rSub of two mice, and in terminal ends of EC projection neurons (outer molecular layers of rDG and rCA2) from one mouse. No mCherry crossover was detected in the left hemisphere. Widespread eGFP expression was detected in lEC and lPaS, as well as lSub, lCA1, and outer molecular layers of lDG. No crossover of eGFP expression into the right hemisphere was detected. ***B***. mCherry and eGFP expression patterns are shown for three individual EC-Tau mice. ***C***. hAPP mice (n=3) exhibit hM4D_i_ DREADDs expression throughout the rMEC, rPaS and rPrS. DREADDs were expressed in the rSub of two mice, while mCherry signal could be seen in the terminal ends of EC projection neurons from one mouse. No crossover of mCherry expression was detected in the left hemisphere. eGFP signal was confined to the left hemisphere, with robust expression in the lMEC, lPaS, lPrS and lSub. eGFP signal was also detected in the outer molecular layers of lDG and in some lDG granule cells. ***D***. mCherry and eGFP expression patterns are shown for all three hAPP mice. ***E***. EC-Tau/hAPP mice (n=4) exhibit hM4D_i_ DREADDs expression throughout the rEC, rPaS and rPrS. No crossover of mCherry expression was detected in the left hemisphere. eGFP expression was less regionally constrained in this group, and appeared in portions of the lMEC, lPaS, and lSub, as well as in the outer molecular layers of lDG, and in lCA1 (stratum radiatum and pyramidale). eGFP expression crossed over into the rPaS of one mouse. ***F***. mCherry and eGFP expression patterns are shown for three individual EC-Tau/hAPP mice. ***G***. mCherry and eGFP expression patterns were combined for all animals (Total, n=12 mice). ***H***. Merged hM4D_i_ DREADDs mCherry expression patterns are shown for 16-month EC-Tau/hAPP mice (n=4) chronically treated with low dose CNO (0.5mg^-1^/kg^-1^/day^-1^). Scale bars, 250µm.

**Supplemental Figure 5.**
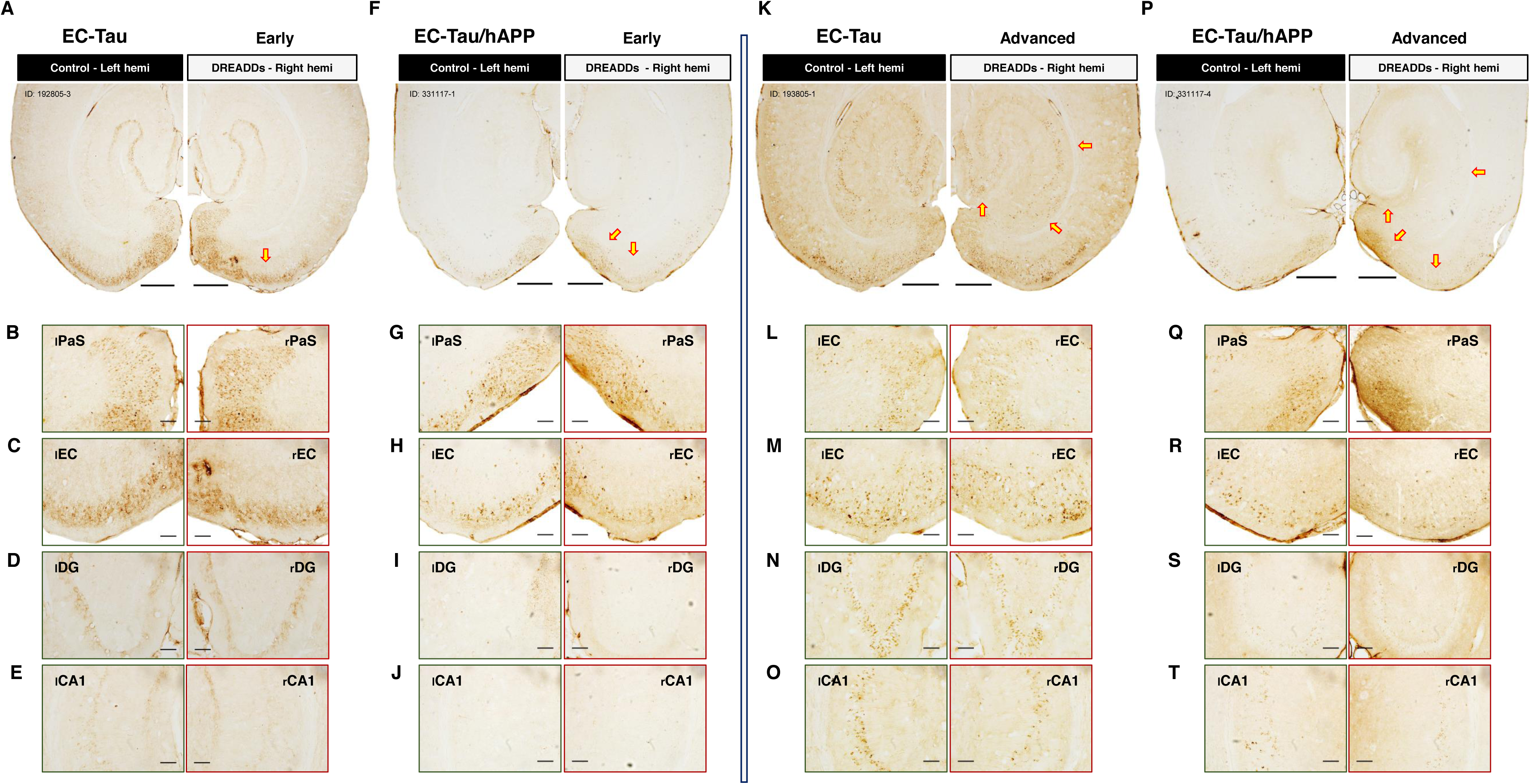
Chronic hM4D_i_ EC DREADDS activation reduces total human tau along the EC-HIPP network. Adjacent tissue sections from hM4D_i_ EC DREADDs activated mice were processed for CP27+ immunostaining experiments to examine the distribution of total human tau expression along the EC-HIPP network. Horizontal brain sections from 16-month EC-Tau/hAPP (n=4) mice and age-matched, littermate EC-Tau (n=5) mice were processed in parallel. ***A.*** EC-Tau, Early Tau pathology; ***F.*** EC-Tau/hAPP, Early Tau pathology; ***K.*** EC-Tau, Advanced Tau pathology; ***P.*** EC-Tau/hAPP, Advanced Tau pathology. Scale bars, 500µm. ***B-E***. An equal distribution pattern of CP27+ immunostaining was detected in PaS cell somas and EC neuropil in an EC-Tau mouse exhibiting Early Tau pathology. ***F-J***. Reduced CP27+ immunostaining was detected in the rPaS and rEC of an EC-Tau/hAPP mouse with Early Tau pathology. ***K-O***. Reduced CP27+ staining was evident in the rDG granule cell layer and rCA1 pyramidal cell layer in an EC-Tau mouse exhibiting Advanced Tau pathology. ***P-T.*** Reduced CP27+ immunostaining was evident in the rPaS, rEC, rDG granule cell layer and rCA1 pyramidal cell layer in an EC-Tau/hAPP mouse with Advanced Tau pathology. Yellow arrows indicate areas of reduced AT8+ immunostaining. All images are representative for human tau-expressing mice that fall into Early Tau pathology and Advanced Tau pathology groupings. Magnified images: scale bars, 100µm.

